# Local translation controls early reactive changes in perisynaptic astrocyte processes at pre-symptomatic stages of Alzheimer’s disease

**DOI:** 10.1101/2025.10.20.683417

**Authors:** Katia Avila-Gutierrez, María Ángeles Carrillo de Sauvage, Marc Oudart, Robin Thompson, Rodrigo Alvear-Perez, Yiannis Poulot-Becq-Giraudon, Esther Kozlowski, Heloïse Monnet, Philippe Mailly, Valentin Garcia, Laurent Jourdren, Alexandre Lucas, Alexis Bemelmans, Helene Hirbec, Carole Escartin, Martine Cohen-Salmon

## Abstract

Early synaptic dysfunction is a hallmark of Alzheimer’s disease (AD), yet the astrocytic mechanisms underlying these alterations remain poorly defined. Here, we identify astrocyte perisynaptic processes (PAPs) as subcellular hotspots of early translational dysregulation in AD. Soluble Aβ_₁–₄₂_ rapidly enhanced global and local protein synthesis in primary astrocytes. In 5.5-month-old APP/PS1-dE9 (APP) mice, translating ribosome affinity purification (TRAP) revealed widespread remodeling of the PAP translatome, while whole-astrocyte translation remained largely unchanged. Dysregulated mRNAs were linked to neuroinflammation, synaptic remodeling, and endoplasmic reticulum stress, and alterations emerged prior to amyloid plaque deposition. Among them, Serpina3n exhibited increased mRNA abundance in PAPs, uncovering spatially restricted translational control. Mechanistically, early Serpina3n upregulation was partially driven by JAK–STAT3 signaling, with preferential effects in astrocyte processes. These findings reveal that local translation in astrocyte PAPs is an early and compartment-specific mechanism that may contribute to synaptic dysfunction and disease initiation in AD.

## Introduction

AD is a neurodegenerative disorder characterized by progressive neuronal loss and decline in cognitive functions. AD is marked by the presence of misfolded protein aggregates, including extracellular Aβ plaques and intracellular neurofibrillary tangles composed of hyperphosphorylated tau [1, 2]. Aβ aggregates spread progressively throughout the brain, with the hippocampus and prefrontal cortex being the first affected regions [3]. The pathological features of AD are diverse and include progressive synaptic dysfunction and neuronal loss, which correlate with the decline in cognitive abilities [4, 5]. Synaptic dysfunction begins with early neuronal hyperactivity, followed by synaptic hypoactivity, and ultimately neuronal degeneration [6, 7]. Understanding the complex mechanisms underlying progressive neuronal loss is fundamental for developing effective therapeutic strategies for this devastating disease.

In recent years, it has become increasingly clear that non-neuronal glial cells, such as astrocytes, microglia, and oligodendrocytes, play crucial roles in neurodegenerative diseases like AD [8]. Astrocytes are vital in maintaining brain homeostasis. They are highly ramified cells and exhibit a high morphological and functional compartmentalization contacting and interacting in a coordinated manner with synapses at the level of PAPs, with brain vessels at the level of perivascular astrocyte processes (PvAPs), and with areas containing the cerebrospinal fluid at the level of the *glia limitans*. In the context of AD, astrocytes undergo a series of functional and morphological changes referred to as reactivity, with both protective and detrimental effects on the brain [9]. Astrocytes are able to counteract the toxic effects of Aβ plaques and tau tangles through various mechanisms, including the clearance of these pathological proteins [10–12]. Reactive astrocytes may also release pro-inflammatory molecules, exacerbating neuroinflammation and promoting neurodegeneration and vascular dysfunction. Importantly, despite being largely overlooked, accumulating evidence indicates that astrocyte dysfunction emerges in the early, presymptomatic phase of AD. Hippocampal astrocyte calcium hyperactivity driven by TRPA1 channel activation is already present in 1-month-old APP mice and coincides with early hippocampal neuronal hyperactivity [13, 14]. Decreased astrocyte calcium activity linked to neuronal hyperactivity is detected in the human cingulate cortex of amyloid accumulators as well as in the 3-month-old *App^NL-F^* mice [15].

Recent studies, including our own, have revealed that locally distributed mRNAs are translated within PAPs and PvAPs [16–18]. This local translation is thought to contribute to the molecular compartmentalization of astrocytes. Importantly, it allows astrocytes to rapidly and locally respond to synaptic changes [19–21]. We also showed altered localization of mRNAs encoding the glial fibrillary acid protein (GFAP) and ferritin in astrocytes in APP mice, an amylosis model of AD [22, 23], suggesting that local translation could be dysregulated in the context of AD.

We here aimed to assess the extent of local translation alterations in astrocytes at the genome wide level, focusing on the pre-symptomatic stage of AD, a period when astrocytic responses remain poorly characterized, despite likely contributing to early synaptic dysfunction [8, 13–15].

## Results

### Soluble Aβ1–42 increases global and local translation in primary astrocytes

Deregulation of local protein synthesis in neurons has been shown to contribute to the development and progression of neurodegenerative diseases such as Alzheimer’s disease (AD) [47–49]. However, very little is known about equivalent defects in astrocytes, which are indispensable partners to neurons in the brain. Here, we first tested the hypothesis that Aβ directly perturbs local translation in astrocytes. Mouse primary cortical astrocytes were exposed to Aβ1–42 oligomers or its diluent DMSO (control) overnight and shortly incubated with puromycin (PMY) (**Fig. 1A**). PMY is an aminoglycoside antibiotic that mimics a charged tRNA^Tyr^ and incorporates in nascent protein chains, allowing visualization of protein synthesis sites [50, 51]. As a negative control, cells were co-incubated with anisomycin to inhibit protein synthesis. Immunodetection of puromycylated chains was performed together with the glial fibrillary acid protein (GFAP) to identify differentiated astrocytes. As expected, PMY signal was abolished in the presence of anisomicyn (**Fig. 1A, C**). In contrast, the signal was higher upon Aβ1–42 treatment compared to DMSO indicating an elevated level of translation (**Fig. 1A, C**). Interestingly, PMY signal appeared also more diffuse and distributed across the entire cells upon Aβ1–42 treatment (**Fig. 1A**). We next repeated these experiments on astrocytes cocultured with neurons, as they display a more differentiated morphology with thin processes (**Fig. 1B**). In these conditions, astrocytes adopted a ramified morphology (**Fig. 1B**). No PMY signal was detected in the presence of anisomycin (**Fig. 1.B, D**). Aβ1–42 treatment resulted in a higher level of PMY signal in astrocyte soma and processes, indicating that translation in both soma and distal processes was stimulated by Aβ1–42 (**Fig. 1B, D**).

**Figure 1.**
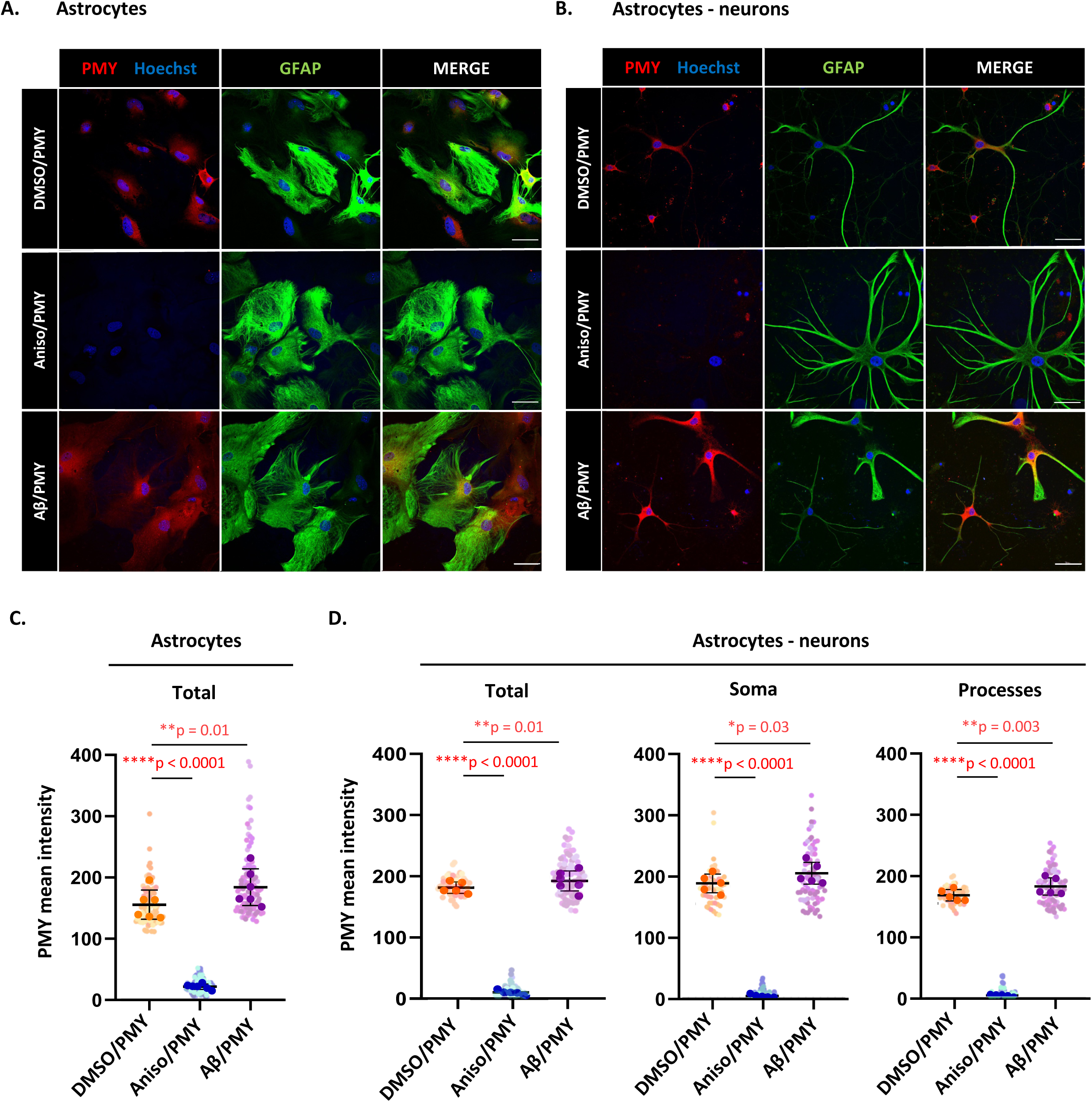
Effect of soluble. **A**β **on local translation in primary astrocytes. (A and B)** Confocal immunofluorescence images showing puromycin (PMY, red) labeling in primary astrocyte cultures **(A)** and astrocyte–neuron co-cultures **(B)**. Astrocytes were treated for 24 h with DMSO or 3 µM soluble Aβ1-42 peptides. They were then incubated 20 min with 3 µM puromycin (PMY) alone or together with 40 µM Anisomycin (Aniso). Astrocytes are immunolabeled with GFAP (green) and nuclei with Hoechst (blue). Scale bar: 20 μm. **(C)** Quantification of PMY mean intensity in whole astrocytes. **(D)** Quantification of PMY mean intensity (signal/area) in astrocyte soma and processes in astrocyte-neurons co-cultures. Data represent individual astrocytes. Mean intensity (signal/area) was calculated for each cell using Fiji’s Measure function. Data were analyzed using a linear mixed-effects model with treatment as a fixed effect and well as a random effect, on Tukey transformed data. Group differences were assessed using Tukey tests. Significant *p*-values (*p* ≤ 0.05) are shown in red. Raw data are provided in **Supplementary Table 5**

These results indicate that Aβ1–42 activates both global and local translation in primary astrocytes.

### Translation is severely altered in hippocampal PAPs in 5.5 months APP mice compared to whole astrocytes

Based on our *in vitro* results, we hypothesized that local translation in PAP is dysregulated in astrocytes in the context of AD. We investigated astrocytic translation in the APP mouse model of AD [24], focusing on the 5.5-month time point—a pre-symptomatic stage when Aβ plaques begin to emerge in the hippocampus [52]. We purified ribosome-bound mRNAs in hippocampal astrocytes and PAPs in WT and APP mice. To do so, we generated an adeno-associated viral (AAV) vector carrying the gfaABC_1_D promoter driving *eGFP-Rpl10a* expression for TRAP experiments [32]. This vector also included MIR124T to prevent potential expression of *eGFP-Rpl10* in neurons [29]. AAV was injected into the hippocampus of 4.5-month-old APP and WT littermate mice as control (**Fig. 2A**). Immunohistochemistry confirmed efficient and specific eGFP-Rpl10a expression in GFAP immunolabeled hippocampal astrocytes (efficiency (infection rate): 87.83% ± 6.02; specificity: 98% ± 0.53) (**Supplementary Fig. 1**). One month after AAV injections, ribosome-bound mRNAs were extracted by TRAP from whole hippocampus (i.e. whole astrocytes) or hippocampal synaptogliosome preparations (i.e. PAPs), the latter comprising pre- and postsynaptic membranes along with PAPs attached to synaptic neuronal membranes [33, 53] (**Fig. 2A**). We confirmed that our RNAseq libraries were enriched in astrocyte-specific mRNAs compared to neuron-, oligodendrocyte-, oligodendrocyte precursor-, mural cell- (vascular smooth muscle cells and pericytes), and endothelial cell-specific mRNAs according to previous previous single cell analyses [54, 55] (**Supplementary Fig. 2**). We then compared WT and APP samples to identify differentially regulated genes (fold-change ≥ 2 adjusted *p* value (*adj-p*) ≤ 0.05) (**Fig. 2B, C; Supplementary Table 1-4**). A comparison of PAPs to whole astrocytes was first carried out to identify PAP-enriched translatomes and their potential changes in APP mice (**Supplementary Table 1, 3, 4)**. In both genotypes, a clear clustering of PAPs and whole astrocyte samples indicated a distinct repertoire of ribosome-bound mRNAs in PAPs compared to astrocytes (**Fig. 2B**). In WT mice, 1,238 genes were enriched and 1,175 were depleted in PAPs compared to whole astrocytes (**Fig. 2B, C; Supplementary Table 1, 4)**. In APP mice, 1,957 genes were enriched and 1,950 were depleted in PAPs compared to astrocytes (**Fig. 2B, C; Supplementary Table 3, 4)**. We then compared APP to WT astrocytes (**Fig. 2B-D; Supplementary Table 4)** and APP to WT PAPs (**Fig. 2B-D Supplementary Table 2, 4)** using the same criteria. Whole astrocyte samples from WT and APP mice were similar (**Fig. 2B, D; Supplementary Table 4**). Only *App* was enriched in APP astrocytes (FC: 2,09; *adj-p* value 3.24 E-5) (**Fig. 2D; Supplementary Table 4**). In great contrast, 167 genes were enriched and 275 were depleted in APP PAPs compared to WT PAPs (**Fig. 2B-D; Supplementary Table 2, 4).**

**Figure 2.**
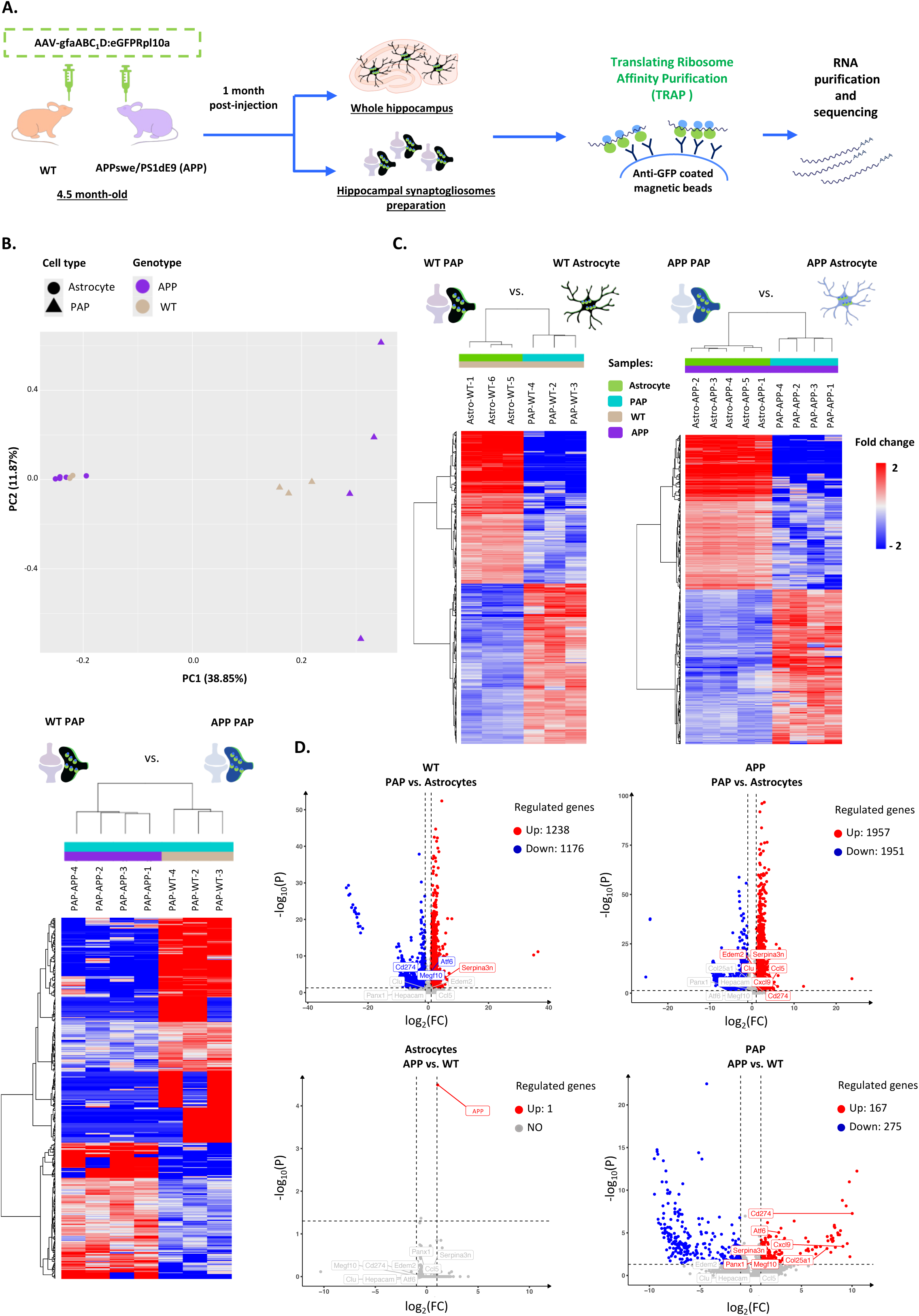
Translation is predominantly altered in perisynaptic processes in 5.5 months APP hippocampal astrocytes. **(A)** Flowchart for the purification of eGFP-Rpl10a astrocyte ribosome-bound mRNAs from whole hippocampi and synaptogliosomes of 5.5-month-old WT and APP mice. (**B)** Principal component analysis of the different libraries showing clear separation of PAP samples from whole astrocytes, with an effect of the APP genotype only on PAP samples. 15.084 genes detected in at least 3 samples were used for this analysis. (**C**) Hierarchical clustering of the differentially expressed ribosome-bound mRNAs in whole astrocytes and PAPs from WT and APP mice. Comparisons are ranked by log_2_ fold change. Red indicates upregulation (log_₂_FC ≥ 1 and padj ≤ 0.05) and blue indicates downregulation (log_₂_FC ≤ −1 and padj ≤ 0.05). (**D)** Volcano plots of the four comparisons between ribosome-bound mRNAs from WT and APP hipoccampal astrocytes and PAPs showing significantly upregulated (log_₂_FC ≥ 1 and padj ≤ 0.05) and downregulated (log_₂_FC ≤ −1 and padj ≤ 0.05) ribosome-bound mRNAs, respectively red and blue dots. Gray dots are mRNAs with unchanged levels (–1< log_₂_FC < 1 or padj > 0.05). Raw data are provided in **Supplementary Table 1-5**.

These findings indicate that, at a pre-symptomatic stage of the disease in APP mice, translation is altered only in PAPs, emphasizing the early specific vulnerability of this astrocyte compartment in AD pathogenesis.

### Local translation changes in APP PAPs involve neuroinflammation, synaptic function and ER stress genes

To identify functional profiles underlying biological processes in our gene sets, we next performed a gene ontology (GO) analysis (**Supplementary Table 1-4**). Several biological processes were over-represented in WT PAPs, when compared to whole astrocytes in WT mice. Most of them were related to neuronal organization such as *neurotransmitter transport* and *axon development* indicating that local translation functionally compartmentalizes astrocytes to support neuronal functions (**Fig. 3A**; **Supplementary Table 1,4**). Interestingly, in APP mice, enrichment scores for these pathways radically changed with a down-representation of *neurotransmitter transport* or *axon development* pathways and an overrepresentation of the *endoplasmic reticulum stress* pathway (**Fig. 3A**; **Supplementary Table 2, 3,4**). Moreover, in APP astrocytes, a huge interferon response was noted, which was also detected in APP PAPs. However, an inverse regulation of *the coagulation cascade*, *inflammation* and *Il6 STAT3 signaling during acute phase response*, which were down represented in APP whole astrocytes but overrepresented in APP PAPs (**Fig. 3B; Supplementary Table 2, 3,4**).

**Figure 3.**
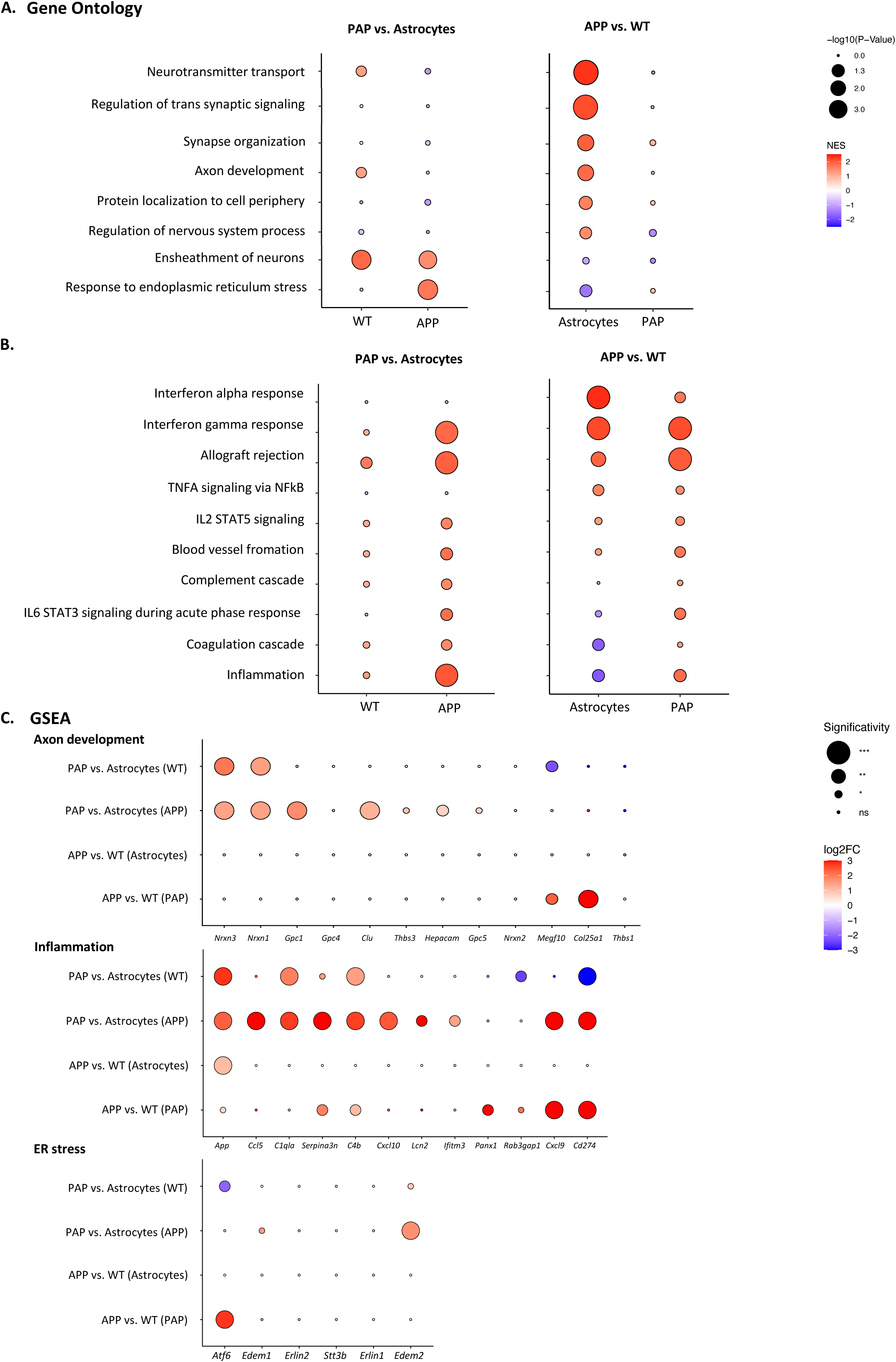
Pathway analysis in 5.5 months APP hippocampal astrocytes and PAPs. **(A)** Gene ontology (GO) enrichment analysis of biological processes in hippocampal APP astrocytes and PAPs for the four comparisons on whole astrocytes or PAPs in WT and APP mice shown in Fig. 1. (**B)** GSEA pathway analysis of the four comparisons. **(A, B)** Dot size indicates –log_₁₀_(*p*-value) and color reflects direction and magnitude of the average fold change. NES, normalized enrichment score. (**C)** Dot plot of the four TRAP-based comparisons showing representative upregulated and downregulated genes associated with specific biological pathways: axon development, inflammation, and ER stress. Dot size indicates significativity, and colors, Log2fold changes (FC). Raw data are provided in **Supplementary Table 1-3**.

We also performed gene set enrichment analyses (GSEA) to determine which sets of genes showed statistically significant changes (**Fig. 3C**). Analyzing up- and down-represented genes in the most significantly changed pathways, we noted, a strong enrichment of several genes previously associated with inflammation and astrocyte reactivity was observed in PAPs from APP mice compared to WT, including *Cd274, Cxcl9, Panx1, Serpina3n and C4b* [9] (**Fig. 3C; Supplementary Table 2, 4**) and in APP PAPs compared to APP astrocytes, including *Cxcl10* and *Ifitm3* [56] (**Fig. 3C; Supplementary Table 3, 4**). We also identified in PAPs from APP mice compared to WT the enrichment of several genes related to synaptic development and remodeling such as *Megf10* [57]and *Col25a1* [58–60] (**Fig. 3C**). Finally, *Atf6*, encoding an important ER stress sensor was highly upregulated in APP PAPs and astrocytes [61, 62] (**Fig. 3C**).

Altogether, these results demonstrate that while inflammatory processes are detectable in APP astrocytes, an increase in translation of genes related to ER stress, neuroinflammation and synaptic function occurs at an early stage in PAPs in young APP mice.

### Levels of mRNAs related to synaptic remodeling, inflammation and ER stress increase in APP synaptogliosomes as early as 3 months

To further characterize the changes occurring in APP PAPs, we analyzed earlier time points to identify the stage at which dysregulation of local translation might be initiated. We focused on genes related to astrocyte reactivity (*Serpina3n, Cd274, Cxcl10 and Ifitm3*), synapse refinement (*Megf10*) and ER stress (*Atf6*). We performed an RT-qPCR analysis on total mRNAs extracted from hippocampal synaptogliosomes obtained from WT or APP aged 2 to 4.5 months (**Fig. 4**). At 2 months, all mRNA levels were similar between WT and APP (**Fig. 4A**). At 3 months, levels of *Serpina3n, Cd274, Megf10, Ifitm3 and Atf6* were higher in APP than WT mice (**Fig. 4A**). Surprisingly, at 4.5 months, we observed more heterogeneity in gene expression and only *Serpina3n* levels remained significantly upregulated in synaptogliosomes from APP mice compared to WT. To further characterized *Serpina3n* upregulation, we performed Western blot assays by capillary electrophoresis to detect Serpina3n in synaptogliosome protein extracts **(Supplementary Fig. 3)**. Unfortunatly, currently used antibodies against Serpina3n [63–65] detected mostly uncharacterized high molecular weight proteins (**Supplementary Fig. 3A, B**) and the two referenced Serpina3n bands around 50 kDa were hardly detectable, although slightly upregulated in APP extracts **(Supplementary Fig. 3C, D).**

**Figure 4.**
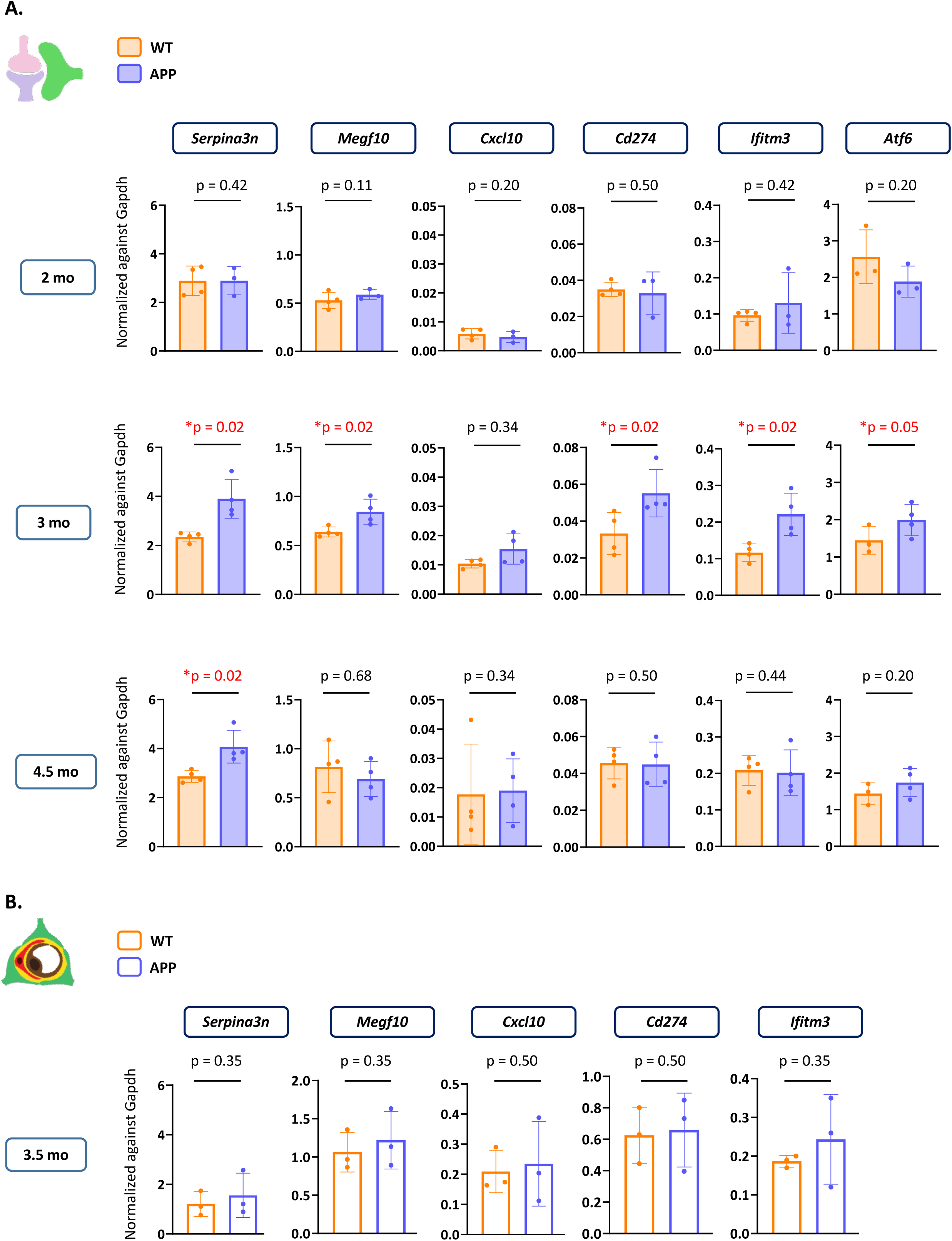
Levels of mRNAs related to synaptic remodeling, inflammation and ER stress in WT and APP PAPs from 3 months. **(A)** qPCR quantification of hippocampal synaptogliosomes in WT and APP mice for *Serpina3n*, *Megf10*, *Cxcl10*, *Cd274*, *Ifitm3*, and *Atf6*. qPCR was performed at 2, 3 and 4.5 months of age. Expression levels were normalized to *Gapdh*. Data are presented as mean ± SD, *n* = 3–4 (one sample per mouse; two hippocampi per sample). (**B)** qPCR quantification of hippocampal purified microvessels (with attached PvAPs) in WT and APP mice for the same genes at 3 months. Expression levels were normalized to *Gapdh*. Data are presented as mean ± SD, *n* = 3 (one sample per mouse; two hippocampi per sample). Statistical analysis: one-tailed Mann–Whitney test. Significant *p*-values (*p* ≤ 0.05) are shown in red; non-significant values are shown in black (*p* > 0.05). Raw data are provided in **Supplementary Table 5**.

To assess whether mRNA changes were affecting other astrocyte subcellular functional domains such as the PvAPs, we also analyzed mechanically purified hippocampal microvessels from 3-month-old mice, on which PvAPs stay attached [31] (**Fig. 4B**). No difference was observed between WT and APP (**Fig. 4B**), suggesting that the changes in early mRNA abundance in AD mice are specific to synaptogliosomes.

These results show that, at the total mRNA level, the upregulation of several genes related to astrocyte reactivity, ER stress and synaptic remodeling is detected in APP synaptogliosomes as early as 3 months.

### *Serpina3n* is upregulated in 5.5 -month-old APP astrocytes soma and processes

To cross-validate our findings on early changes in PAP-translated mRNA in APP mice, we performed FISH on 5.5 -month-old hippocampal sections. This allowed the direct detection of *Serpina3n* mRNAs, which encodes a serine protease inhibitor that plays a critical role in reactive astrocytes [63, 64, 66], and analysis of its subcellular localization within GFAP labelled astrocytes (**Fig.** 5**A**). Each astrocyte was delineated manually in 3D on the basis of GFAP immunolabeling [22] (**Fig. 5A**). *Serpina3n* mRNAs were highly associated to astrocytes (**Fig. 5A**) and not to microglia stained by Iba1 (**Supplementary Fig. 4A**). The number of *Serpina3n* FISH dots in the soma or processes was calculated for each astrocyte and normalized to the cell volume (**Fig. 5B**). Importantly, we considered astrocytes not in direct contact with Methoxy-X04^+^ amyloid plaques, which started to develop in some mice. Astrocyte volumes were not increased in APP compared to WT mice (**Fig. 5A, B**). However, GFAP+ main processes were enlarged in APP as compared to WT astrocytes, a typical reactive morphological change (**Fig. 5A**). The number of *Serpina3n* FISH dots in both astrocyte soma and processes was higher in APP mice (**Fig. 5B**). At 5.5 month, few amyloid plaques started to be detected in half of the analysed APP mice. Thus, we analysed *Serpina3n* by FISH specifically in astrocytes proximal to amyloid plaques (**Supplementary Fig. 5**). Compared to astrocytes distant form amyloid plaques, these astrocytes tended to show larger volumes, and higher levels of *Serpina3n* mRNA in their soma and processes (**Supplementary Fig. 5**), indicating that amyloid plaques might have a direct effect in the expression of *Serpina3n* in astrocytes at early stages.

**Fig. 5.**
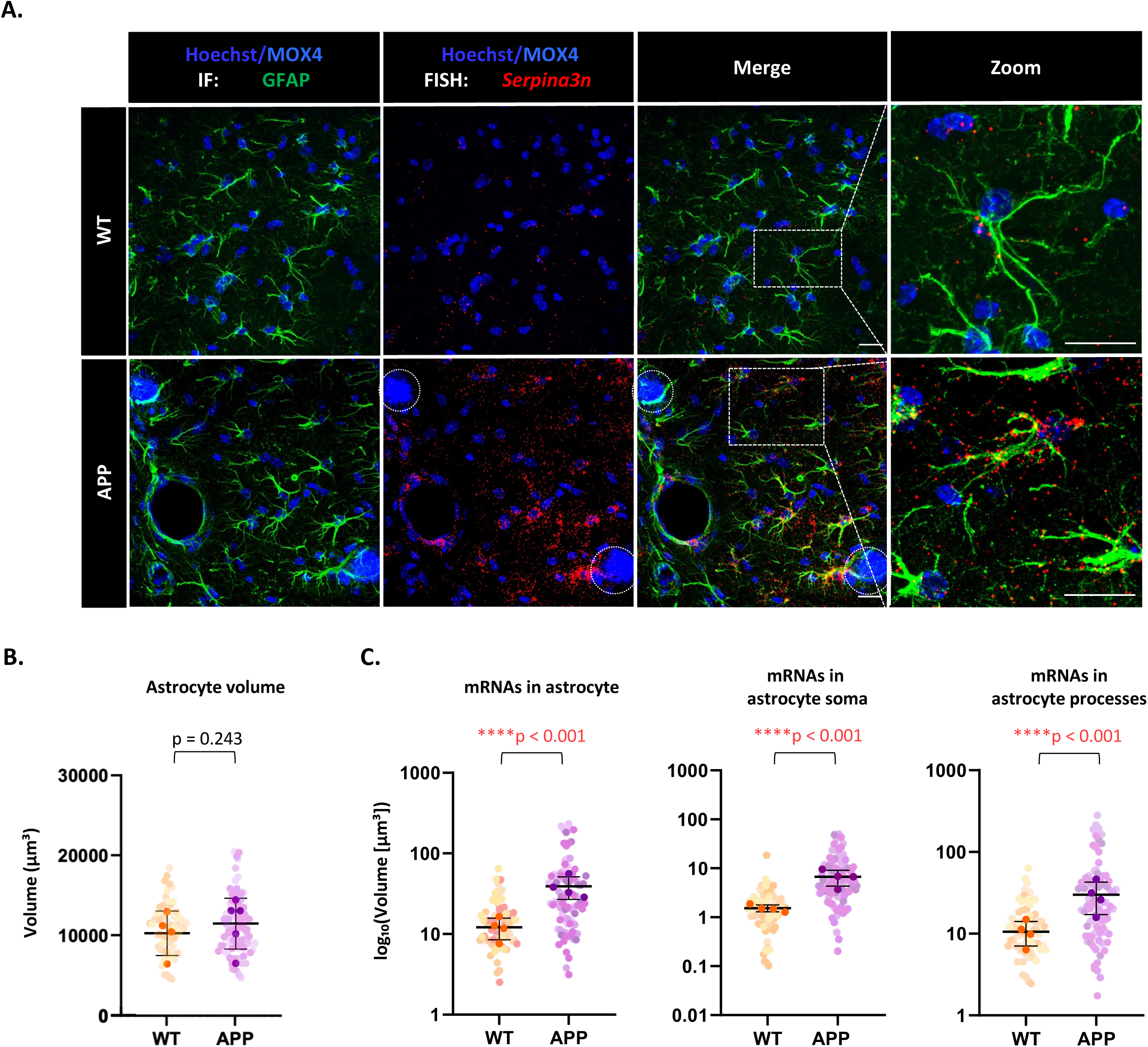
*In situ* characterization of *Serpina3n* mRNAs in 5.5 months APP astrocytes. **(A)** Deconvoluted confocal microscopy images showing FISH detection of *Serpina3n* mRNAs (red dots) in CA1 hippocampal astrocytes immunolabeled with GFAP (green). Amyloid-β (Aβ) plaques were stained with Methoxy-X04, and nuclei were counterstained with Hoechst (blue). Dotted circles outline Aβ plaques. Scale bar: 20 µm. (**B-C)** FISH quantification in CA1 hippocampal astrocytes from WT and APP/PS1 mice: (**B**) astrocyte volume, (**C**) total *Serpina3n* mRNA density in whole astrocytes, soma, and processes. FISH dot density was normalized to individual ROI volumes. Data are presented as mean ± SD. N = 4 mice (4 images per mouse, 70 individual astrocytes in total) WT and N=4 mice (4 images per mouse, 87 astrocytes) APP astrocytes were analyzed. Data were analyzed using a linear mixed-effects model with genotype as a fixed effect and mouse as a random effect on Tukey transformed data. Comparison between the two groups were performed using T-test. Significant *p*-values (*p* ≤ 0.05) are shown in red; non-significant values are shown in black. Raw data are provided in **Supplementary Table 5**

These results show that, in contrast to ribosome-bound mRNAs, the level of *Serpina3n* total mRNA is globally higher in APP astrocytes (soma and processes) at 5.5 month compared to WT mice. Thus, specific local mechanisms occur to activate translation of *Serpina3n* in PAPs.

### *Serpina3n* upregulation in APP astrocytes is detected at 3.5 month and involves the JAK/STAT3 pathway

We next investigated the mechanisms that might drive *Serpina3n* upregulation in astrocytes and its localization within PAPs. Our previous RT-qPCR analysis (**Fig. 4**) revealed increased *Serpina3n* mRNA levels in synaptogliosomes from 3 months, suggesting this time point marks the onset of *Serpina3n mRNA* accumulation in PAPs. We focused on the JAK-STAT3 pathway, a central cascade controlling astrocyte reactivity [67] which was over-represented in APP PAPs (**Fig. 3**). Activation of the JAK-STAT3 pathway in hippocampal APP astrocytes was previously shown to induce astrocyte reactivity and upregulate *Serpina3n* and its inhibition by the inhibitor ‘Suppressor Of Cytokine Signaling 3’ (SOCS3) to downregulate *Serpina3n* [29]. We first performed FISH on 3.5-month-old hippocampal slices (**Fig. 6**). As done previously (**Fig. 5**), each individual astrocyte domain was delineated in 3D on the basis of GFAP immunolabeling (**Fig. 6B**). The astrocyte volume was unchanged between WT and APP (**Fig. 6B, C**). Very few FISH dots were detected in WT (**Fig. 6B, D**). In great contrast, in APP, *Serpina3n* mRNA dots were numerous in astrocyte soma and processes (**Fig. 6B, D-E**). Interestingly, a high variability between individuals was observed, indicating that the 3.5-month stage may represent a transitional phase during which astrocyte reactivity begin to emerge. In addition, as observed at 5.5 months, the level of *Serpina3n* total mRNA was globally higher in APP astrocytes (soma and processes).

**Fig. 6.**
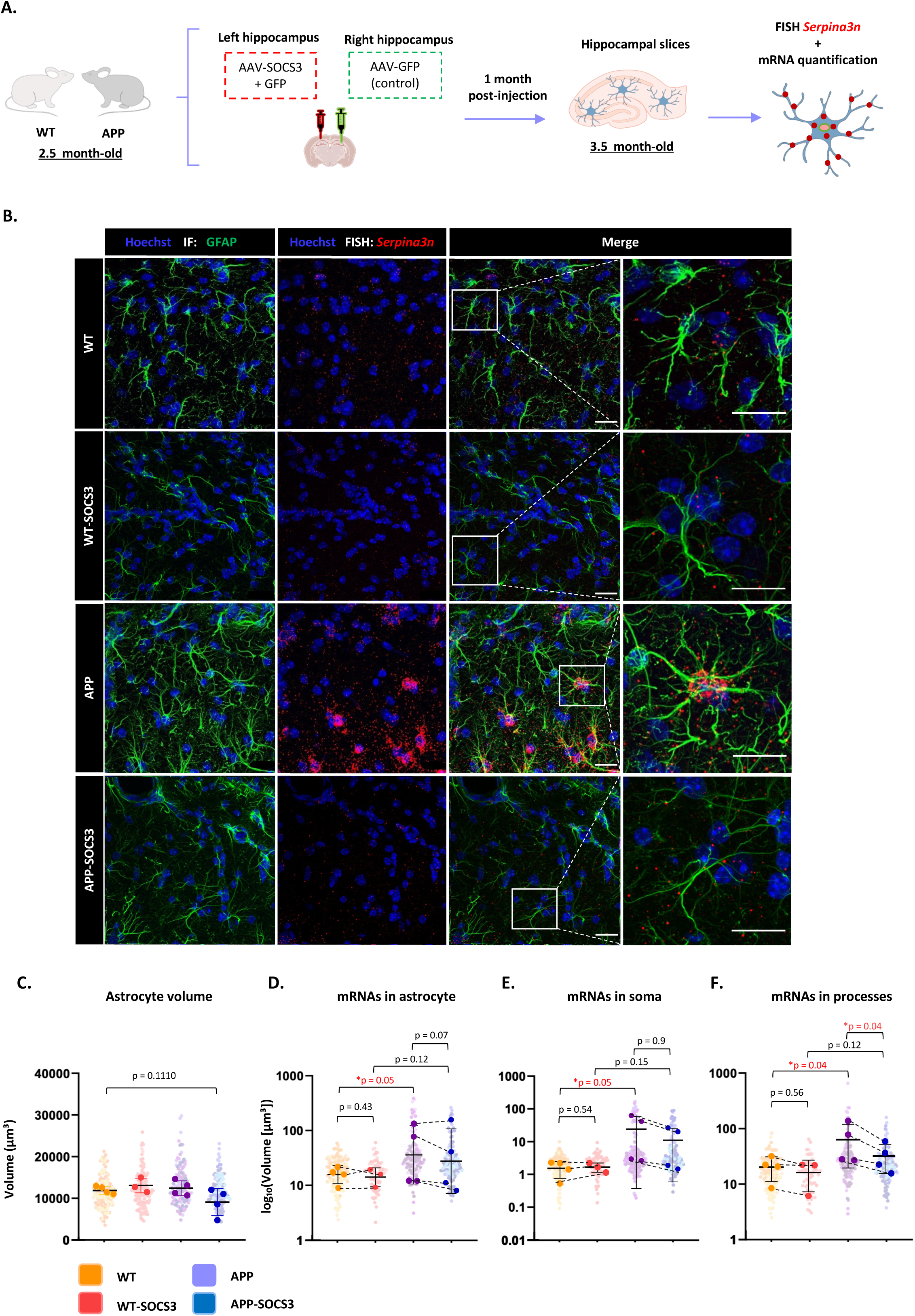
*In situ* characterization of *Serpina3n* mRNAs in 3.5 months APP astrocytes. Involvement of the JAK/STAT3 pathway. **(A)** Flowchart of the experimental procedure following bilateral hippocampal injections in 2.5-month-old WT and APP mice. The right hemisphere was injected with AAV-GFP, and the left with AAV-SOCS3+GFP. One month later, brains were fixed by PFA perfusion, dissected and analyzed. (**B)** Deconvoluted confocal microscopy images showing FISH detection of *Serpina3n* mRNAs (red dots) in CA1 hippocampal astrocytes immunolabeled with GFAP (green). Nuclei were counterstained with Hoechst (blue). Scale bar: 20 µm. (**C-E)** FISH quantification in CA1 hippocampal astrocytes from WT and APP mice: (**C**) astrocyte volume, (**D**) total *Serpina3n* mRNA density in whole astrocytes, (**E**) soma, and (**F**) processes. FISH dot density was normalized to individual ROI volumes. Data are presented as mean ± SD. Sample sizes were as follows: WT-GFP, n = 4 mice (4 images per mouse, 171 individual astrocytes in total); WT-SOCS3, n = 3 mice (4 images per mouse, 68 astrocytes); APP-GFP, n = 4 mice (4 images per mouse, 128 astrocytes); and APP-SOCS3, n = 4 mice (4 images per mouse, 124 astrocytes). Data were analyzed using a linear mixed-effects model with treatment and genotype, as well as their interaction (treatment × genotype), as fixed effects, and mouse as a random effect. Tukey transformation was used for whole astrocytes and soma, and log transformation for astrocytic processes). Post hoc multiple comparisons were performed using Tukey’s tests. Significant *p*-values (*p* ≤ 0.05) are shown in red; non-significant values are shown in black (*p* > 0.05). Raw data are provided in **Supplementary Table 5**.

We next inhibited the JAK-STAT3 pathway by overexpressing SOCS3 in 2.5 months APP mice. We used AAV that target astrocytes to express SOCS3 or eGFP in CA1 hippocampal astrocytes [29] (**Fig. 7A**). AAV-SOCS3 was co-injected with AAV-eGFP in one hippocampus to visualize infected astrocytes in APP (APP-SOCS3) and WT (WT-SOCS3) mice. Controls were the same mice injected in the other hippocampus with AAV-GFP alone, at the same total viral titer (APP-GFP and WT-GFP respectively). Immunohistochemistry confirmed efficient and specific *eGFP* expression in GFAP immunolabeled hippocampal astrocytes (efficiency (infection rate) 91.06% ± 4.18; specificity 99% ± 2.45) (**Supplementary Fig. 6**). The same variability in *Serpina3n* mRNA level was observed for APP-SOCS3 compared to APP. However, compared to APP, *Serpina3n* mRNA levels tended to decrease in whole astrocytes and significant difference with WT was lost (**Fig. 6B, D**). The same was observed in the soma (**Fig. 6E**). However, in processes, we observed a significant rescue of the mRNA levels (**Fig. 6E**) suggesting that the inhibitory effect of SOCS3 was more pronounced in astrocyte processes.

Overall, these results indicate that levels of mRNAs in astrocyte soma and processes is already increased on 3.5 months in APP mice. The inactivation of the JAK-STAT3 in astrocytes is able to partially reduce the expression of *Serpina3n* in astrocyte processes specifically.

## Discussion

Our study proposes that alterations in the molecular repertoire of PAPs is driven by local translation and is an early pathogenic mechanism in APP mice. Changes concerned genes involved in astrocyte reactivity, ER stress, neuronal development, and synaptic function. At the level of total mRNAs upregulation was observed in PAPs, but not in PvAPs, from 3 months, a stage when no amyloid plaques are yet detected. Focusing on *Serpina3n*, a marker of astrocyte reactivity, we found that while increase in translation was restricted to PAPs, total mRNA was also upregulated in the astrocyte soma, suggesting the activation of translation mechanisms specifically in PAPs. Of note, soluble Aβ applied on primary astrocytes increased global as well as local translation. Finally, we showed that the level *Serpina3n* total mRNAs in PAPs was sensitive to the JAK-STAT3 pathway activity. Collectively, our data suggest that early local translation changes in astrocytes occur in PAPs in APP mice and may contribute to astrocyte reactivity and dysregulated synaptic homeostasis that may drive disease progression.

Serpina3n belongs to the serpin family, which includes serine protease inhibitors and acute-phase proteins [68]. Within the central nervous system, astrocytes are the primary source of Serpina3n [63], which is predominantly secreted into the extracellular space [64]. Serpina3n upregulation is a canonical marker of astrocyte reactivity [9]. Several studies reported increased levels of the human homolog *SERPINA3* in specific brain regions of AD aged patients, including the hippocampus and frontal cortex [69]. Elevated intracellular and extracellular levels of Serpina3n has been observed near plaques in both AD patients and transgenic mouse models [70]. It has been proposed that SERPINA3 acts as a pathological chaperone, colocalizing with plaques and promoting Aβ aggregation [70]. Structural studies suggest that Aβ specifically interacts with two β-sheets of SERPINA3, potentially stabilizing plaques and rendering them more resistant to degradation, thereby exacerbating neuronal damage [71]. A bioinformatic analysis recently suggested the correlation between *SERPINA3N* upregulation and astrocyte markers in elderly AD patients [69]. Interestingly, we observed that in APP astrocytes proximal to newly formed amyloid plaques, the level of *Serpina3n* tend to be higher but was not dramatically different from more distal astrocytes. Thus, there is not a clear correlation between the presence of Aβ aggregation and expression of *Serpina3n* in astrocytes. More surprisingly, an increased astrocyte expression of *Serpina3n* was observed as early as 3 months, before hippocampal amyloid plaque formation. Unfortunately, due to the low specificity of the most currently used Serpina3n antibodies and the probably low amount of Serpina3n in synaptogliosomes, we could not observe if elevation of total *Serpina3n* mRNA in APP was reflected at the protein level. Altogether, previous studies our data suggest that elevation of Serpina3n in astrocytes and particularly at their interface with neurons is marker presymptomatic marker of AD that may contribute to initiate Aβ aggregation.

The JAK-STAT3 pathway is a key regulator of astrocyte reactivity in various neurodegenerative diseases, including Huntington’s disease and AD [67]. Inhibition of this pathway via astrocyte-specific overexpression of SOCS3, which blocks the interaction between STAT proteins and JAK [72, 73], has been shown to prevent astrocyte reactivity and upregulation of total mRNAs such as *Serpina3n* [29]. We showed here that inhibition of the JAK-STAT3 pathway by SOCS3 at a very early stage of AD reduces Serpina3n levels specifically in PAPs. Thus, the activation of JAK-STAT3 seems to occur at very early stages of AD and seems to potentiate the distribution of *Serpina3n* mRNAs in PAPs. SOCS3 inhibits the expression of several mRNAs typical of astrocyte reactivity in APP astrocytes [9]. Thus, as shown for *Serpina3n*, it might also influence their distribution and translation in PAPs. To fully answer this question, a future TRAP RNAseq comparison could be performed between APP and APP-SOCS3 PAPs. Interestingly, STAT3 has also non-canonical activities and was recently identified as an RNA-binding protein [74, 75]. It would therefore be compelling to investigate whether STAT3 directly regulates mRNA stability in astrocytes.

An obvious question raised by our results concerns the mechanisms underlying local translation regulations in PAPs. The comparison of *Serpina3n* expression using FISH, RT-qPCR, and ribosome-bound mRNA sequencing provided valuable insights. It revealed that total mRNA levels were upregulated in both astrocyte soma and processes, whereas ribosome-bound mRNAs were upregulated specifically in PAPs. This dichotomy suggests different possible scenarios: 1) mRNAs free of ribosomes are transported in PAPs and bind to ribosomes only on site; 2) Ribosome-bound mRNAs compacted in granules but inaccessible to TRAP are transported and released in PAPs for local translation. In both hypotheses, soluble Aβ1–42 peptides may act as the trigger, since they modified translation in primary astrocytes, as well as local translation in their processes. Synaptic alterations may signal locally to astrocytes as they have been found as early as 3 months in other AD mouse models [13, 14]. Moreover, previous studies, including our own, demonstrated that local translation in PAPs is plastic and responsive to changes in neurotransmission [19, 20]. It would therefore be interesting to mechanistically address the specific role of AD-linked early synaptic alterations on local translation in astrocytes. Interestingly, the same mechanisms would not occur at the vascular levels, since at least for the tested mRNAs, no upregulation could be found in purified gliovascular units.

Aside *Serpina3n*, several other canonical markers of astrocyte reactivity were upregulated specifically in APP/WT PAPs or APP PAPs/Astrocytes [76]. The interferon-induced transmembrane protein 3, Ifit3m, upregulated in APP PAPs/astrocytes, was recently shown to bind and upregulated γ-secretase activity increasing the production of Aβ [56]. *Cd274* encoding Pd-l1 has been shown to reduce chronic neuroinflammation in the context of multiple sclerosis [77], and in APP mice to sustain microglial Aβ uptake [78]. They could have opposite effects in APP PAPs.

Until recently, altered astrocyte reactivity was associated with late AD, i.e. when amyloid plaques are already developed. Our findings indicate that, already at early stages of AD in the APP mouse model, profound alterations in local translation occur within hippocampal PAPs. These changes are associated with astrocyte reactivity, ER stress, and synaptic remodeling. Activation of the JAK–STAT3 signaling pathway at these early stages may initiate them. Thus, in the context of AD, PAPs emerge as pathological hotspots where early changes in mRNA localization and local translation may both sense and contribute to synaptic dysfunction, preceding widespread neuronal degeneration.

## Methods

### Mice

APP transgenic mice on the C57BL6J background (https://www.jax.org/strain/005864) were used in this study [24]. They are double transgenic mice expressing a chimeric mouse/human amyloid precursor protein (Mo/HuAPP695swe) and a mutant human presenilin 1 lacking exon 9 (PS1-dE9). Both mutations are associated with early-onset AD. Non-transgenic littermates were used as controls. All experimental protocols were reviewed and approved by the local ethics committees (CETEA #44 and CEEA59) and submitted to the French Ministry of Education and Research (Approvals # APAFIS4565,4503, 38200 and 4565-20). They were performed in strict accordance with recommendations of the European Union (2010–63/ EEC).

TRAP experiments, qPCR on purified gliovascular unit and FISH analyses were all performed on males. qPCR on synaptogliosomes were performed on females. This choice was driven by our commitment to utilize all animals born for our experiments.

### In vitro study

#### Primary astrocytes

To obtain astrocyte-enriched cultures, postnatal day 1 to 4 pups were used. Before starting, astrocyte growth medium was prewarmed to 37 °C (DMEM supplemented with 10% heat-inactivated fetal bovine serum [hiFBS], 1% penicillin/streptomycin, and 1× GlutaMAX). Cell culture flasks were coated with poly-D-lysine (PDL, 0.1 mg/mL) at 37 °C for at least 2 h and then rinsed twice with PBS. All cultures were maintained at 37 °C in a 5% CO_₂_ incubator. Each pup’s head was well rinsed with 70% ethanol and sacrificed by decapitation. The brain was placed in ice-cold PBS in a dissection dish. The olfactory bulbs and cerebellum were removed, and the remaining tissue was mechanically dissociated into single cells by suspending in culture medium and triturating 10–15 times with a glass Pasteur pipette. The cell suspension was transferred to a 15 mL conical tube, brought to 15 mL with PBS, and centrifuged at 100 rcf for 5 min. The pellet was resuspended in 10 mL of astrocyte medium and plated onto PDL-coated flasks. Medium was replaced every 3 days until cultures reached 70–80% confluency. To reduce microglial contamination, cultures were shaken at 180 rpm for 4 h at 37 °C on an orbital shaker, and the medium was replaced immediately afterwards. Cells were then trypsinized, and 50,000 cells were seeded into each well of a PDL-coated 24-well plate containing a glass coverslip. After 24 h, the medium was replaced with reduced-serum astrocyte medium (DMEM, 5% hiFBS, 1% penicillin/streptomycin, 1× GlutaMAX) supplemented with 4 µM cytarabine (AraC) to limit proliferation. Cultures were maintained in this medium for two weeks with medium changes every 3 days.

#### Co-culture of astrocytes and hippocampal neurons

The co-culture model was established as described by Luchena et al. [25], with minor modifications. Astrocytes were prepared as outlined above; however, after homogenization, half of a whole brain was divided equally into 24 tubes, and each aliquot was plated directly into a PDL-coated 24-well plate containing a glass coverslip in astrocyte growth medium. After 72 h, a confluent astrocytic monolayer was formed in each well, and the medium was replaced with neuronal plating medium (Neurobasal [Thermo] supplemented with 10% hiFBS, 1% penicillin/streptomycin, and 1× GlutaMAX). Hippocampal neurons were prepared from embryonic day 18 brains. The hippocampus was dissected out. After dissociation and centrifugation, cells were resuspended in neuronal plating medium, and viable cells were quantified using trypan blue to identify dead cells. A total of 50,000 live neurons were seeded onto the astrocytic monolayer in each well. After 24 h, the medium was replaced with neuronal growth medium (Neurobasal supplemented with 1× B27, 1% penicillin/streptomycin, and 1× GlutaMAX) containing the antimitotic agents 5-fluoro-2’-deoxyuridine and uridine (FDU+U) at a final concentration of 20 µM to inhibit glial proliferation. Medium was renewed every 2 days to maintain FDU+U levels. After 10 days in FDU+U-supplemented neuronal growth medium, co-cultures were ready for further processing and experimental treatment.

#### Amyloid-β (Aβ1–42) exposure

Oligomeric Aβ1–42 was prepared as reported previously [26, 27]. Aβ1–42 was dissolved in 100% hexafluoroisopropanol (HFIP) to a concentration of 1 mM. HFIP was then evaporated under vacuum, and the resulting peptide film was stored at −20 °C until use. For oligomer preparation, the peptide was resuspended in dry dimethylsulfoxide (DMSO, 5 mM) and subsequently diluted in Ham’s F-12 pH 7.4 culture medium to a final concentration of 100 μM, followed by incubation at 4 °C for 24 h. Before cell treatment, the Aβ1–42 solution was diluted in growth medium to an intermediate concentration of 30 μM. Cells were then exposed to Aβ1–42 at a final concentration of 3 μM for 24 h at 37 °C in a 5% CO2 humidified incubator. DMSO was used as vehicle control.

#### Puromycylation

Cells were incubated with 3 μM puromycin (PMY) diluted in culture medium for 20 min at 37 °C. DMSO was used as vehicle control. To inhibit protein synthesis, cells were pretreated with 40 μM anisomycin (dissolved in DMSO) for 20 min at 37 °C, prior to puromycin exposure. Cells were then washed twice with PBS and subsequently fixed in 4% paraformaldehyde containing 4% sucrose in PBS for 20 min at room temperature. Cells were then rinsed for 5 min in PBS three times and permeabilized and blocked in 3% BSA, 100 mM glycine, and 0.25% Triton X-100 for 1 h. Next, slices were incubated overnight at 4°C with primary antibodies in the blocking solution. Then, cells were rinsed 3 times in PBS for 10 min, and incubated with secondary antibodies for 2 h at room temperature. Cells were rinsed again 3 times with PBS and mounted in Fluoromount (Southern Biotech, Birmingham, AL) on glass slides. The antibodies and probes are listed in the **Supplementary Table 6.** Images were acquired with a Zeiss spinning-disk confocal microscope using either a 40× or 63× objective, collecting z-stacks of 4 planes with a 0.3 μm step size.

A quantitative analysis of PMY immunolabeling was performed by measuring mean fluorescence intensity from immunofluorescence images. Regions of interest (ROIs) were individual astrocytes. Cells limits were defined based on glial fibrillary acid protein (GFAP) immunolabeling. The soma was delineated by selecting the Hoechst-stained nucleus plus 2 μm to include the perinuclear cytoplasm. Puromycin intensity was then obtained using the *Measure* function in Fiji, and values were normalized to the area of the corresponding ROI.

### Viral Vectors

We used adeno-associated virus (AAV2, serotype 9) that bear the gfaABC_1_D promoter, a synthetic promoter derived from the GFAP promoter [28], to drive expression of eGFP-Rpl10a (TRAP gene) in astrocytes and 4 copies of the *MiR124* target sequence to block transgene expression in neurons [29]). AAV were produced by the MIRCen viral vector facility according to validated procedures [30]. Viral genome concentration in the vector batch was determined by qPCR on DNase resistant particles. To inhibit the JAK2-STAT3 pathway in mouse astrocytes, we used an AAV encoding murine *Socs3* under the same gfaABC1D promoter and 4 copies of the *MiR124* target sequence as described previously [29]. Control viral vectors encoded *Gfp*. AAV-SOCS3 were co-injected with an AAV-GFP to visualize infected cells (same total viral titer).

### Stereotaxic injections

Mice were anesthetized with an intra peritoneal injection of ketamine (100 mg/kg) and medetomidine (0.25 mg/kg) and anesthesia was reversed by a sub cutaneous injection of atipamezole (0.25 mg/kg) at the end of the surgical procedure. Mice were given a sub cutaneous injection of buprenorphine (0.075 mg/kg) and xylocaine (5 mg/kg, at the incision site) 5 min prior to surgery. Mouse temperature was maintained at 37°C throughout the procedure, thanks to a heating pad connected to a rectal temperature probe. Mice were given sub cutaneous injection of 100 µl of warm saline before and after surgery to prevent dehydration.

Viral vectors were injected in the CA1 region of the hippocampus (coordinates from Bregma: anteroposterior (AP): − 3, lateral (L): +/− 3; ventral (V): − 1.5 mm from the dura). AAV were diluted in 0.1 M phosphate buffer saline (PBS) with 0.001% pluronic acid, at a final total concentration of 2.5 10^9^ viral genome (VG)/μl. Two μl of viral suspensions were injected at a rate of 0.25 μl/min. AAV-eGFP-RPL10 was injected bilaterally, while AAV-GFP and AAV-SOCS3+GFP were injected in the left and right hippocampus of the same mouse respectively.

### Synaptogliosome preparation

Synaptogliosomes were prepared from whole hippocampi as previously described [18]. All steps were performed at 4°C. Fresh dissected hippocampi were homogenized with a tight glass homogenizer (20 strokes) in buffer solution (0.32 M sucrose and 10 mM HEPES in diethyl-dicarbonate-treated water, with 0.5 mM DTT, Protease Inhibitors (Complete-EDTA free) 1 mini tablet/10 mL, RNasin1μL/mL, Cycloheximide (CHX) 100 μg/mL added extra-temporaneously). The homogenate was then centrifuged at 900 g for 15 min. The pellet was discarded, and the supernatant was centrifuged at 16,000g for 15 min. The second supernatant was discarded, and the second pellet (containing synaptogliosomes) was diluted in 600 µl of buffer solution and centrifuged again at 16,000 g for 15 min. The final pellet contained the synaptogliosomes. They were stored at −80°C for RNA preparation or immediately processed for TRAP.

### Preparation of microvessels

This procedure allows to purify brain microvessels on which perivascular astrocyte processes (PvPAs) remain attached. Fresh dissected hippocampi were resuspended in HBSS/HEPES using an automated douncer at 4°C. After a first centrifugation at 2000 g for 10 min, the pellet was resuspended in HBSS/Dextran 18% and centrifuged at 4000 g for 15 min to separate myelin from brain vessels. This new pellet containing brain vessels with PvAPs attached was resuspended in HBSS/BSA 1%. Filtration on a 20 µm-mesh filter allowed brain microvessels retention. This protocol is described in full details in [31].

### Translating ribosome affinity purification (TRAP)

TRAP is an immunopurification of eGFP-tagged ribosomes [32]. Extractions were performed on fresh dissected whole hippocampi or synaptogliosomes of AAV-gfaABC_1_D:eGFP-Rpl10a injected mice (one month after injection) following our refined TRAP protocol described in full details in [33]. For RNAseq libraries, we pooled 2 hippocampi of the same mouse or synaptogliosomes prepared from 4 hippocampi from 2 mice. Eluted ribosome-bound mRNAs were purified using the RNeasy Lipid tissue kit (Qiagen) and stored at −80°C.

### RNA-Seq and analysis

Ten nanograms of total RNA were amplified and converted into cDNA using a SMART-Seq v4 Ultra Low Input RNA kit (Clontech). Next, an average of 150 pg of amplified cDNA was used per library, with a Nextera XT DNA kit (Illumina). Libraries were multiplexed on two high-output flow cells and sequenced (75 bp reads) on a NextSeq 500 device (Illumina). The mean ± standard deviation number of reads per sample meeting the Illumina quality criterion was 23 ± 6 million.

Analysis was performed by GenoSplice technology (Paris, France). Analysis of sequencing data quality, reads repartition (*e.g.*, for potential ribosomal contamination), inner distance size estimation, genebody coverage, strand-specificity of library were performed using FastQC v0.11.2, Picard-Tools v1.119, Samtools v1.0, and RSeQC v2.3.9. Reads were mapped using STAR v2.7.5a [34] on the mouse mm39 genome assembly and read count was performed using featureCount from SubRead v1.5.0 and the Human FAST DB v2022_1 annotation. Sample genotype was confirmed by manually inspection of BAM using IGV (presence of the Swedish mutations in human *APP* and skipping of the human *PSEN1* exon 9 in APP samples). The RNA-seq gene expression data and raw fastq files are available on the GEO repository (www.ncbi.nlm.nih.gov/geo/) under the accession number GSE307321.

Gene expression was estimated as described previously [35]. Only genes expressed in at least one of the two compared conditions and covered by enough uniquely mapped reads were further analyzed. Genes were considered as expressed if their FPKM value was greater than FPKM of 98% of the intergenic regions (background). At least 50% of uniquely mapped reads was required. Analysis at the gene level was performed using DESeq2 [36]. Genes were considered differentially expressed for fold-changes ≥ 2 and p-adjusted (adj) values ≤ 0.05. Pathway enrichment analyses and Gene Set Enrichment Analysis

(GSEA) analysis [37] were performed using WebGestalt v0.4.4 [38] merging results from up-regulated and down-regulated genes only, as well as all regulated genes. Pathways and networks were considered significant with p-adjvalues ≤ 0.05. GSVA on gene sets of interest was performed using R [39].

### FISH and GFAP co-immunofluorescent detection on mouse brain sections

Fixed hippocampal slices (40 µm thick) were carefully washed three times with 1× PBS in a 24-well plate. During the final wash, the PBS was replaced with 4 drops of RNAscope® Hydrogen Peroxide Solution (Advanced Cell Diagnostics Inc.) and incubated for 10 min at room temperature (RT) to block endogenous peroxidase activity, which resulted in the appearance of small bubbles. The slices were then rinsed with Tris-buffered saline containing Tween® 20 (50 mM Tris-Cl, pH 7.6; 150 mM NaCl; 0.1% Tween® 20) at RT and mounted onto Super Frost+®-treated glass slides using a paintbrush.

The mounted slices were air-dried at RT for 1 h in the dark, briefly immersed (less than 3 s) in deionized water in a glass chamber, and air-dried again for 1 h at RT in the dark. They were then incubated for 1 h at 60°C in a dry oven, followed by overnight drying at RT in the dark. For rehydration, slices were quickly submerged (less than 3 s) in deionized water at RT. Excess liquid was blotted with absorbent paper, and a hydrophobic barrier was drawn around the tissue. A drop of pure ethanol was applied to the slice for less than 3 s and promptly removed using absorbent paper. The slides were then incubated in a steamer at 100°C. A drop of preheated RNAscope® 1× Target Retrieval Reagent (Advanced Cell Diagnostics Inc.) was added to the steamer, and the slides were incubated for 15 min. Then, the slides were washed three times in deionized water at RT, and excess moisture was removed. A drop of 100% ethanol was applied for 3 min and then blotted. Next, a drop of RNAscope® Protease+ solution (Advanced Cell Diagnostics Inc.) was added, and the slices were incubated at 40°C for 30 min in a humid chamber. The target retrieval and Protease+ treatments were used to unmask mRNA targets. FISH was performed according to the v2 Multiplex RNAscope technique (Advanced Cell Diagnostics, Inc., Newark, CA, USA). After the FISH procedure, GFAP was detected by immunofluorescence. Independent experiments were performed on 4 animals per genotype and 3 brain slices per animal. This protocol is detailed in [40, 41]. Images were acquired using a Yokogawa W1 Spinning Disk confocal microscope (Zeiss) with a a 63X oil objective and the Metamorph Premier 7.8 software. The FISH and GFAP channel were deconvoluted with Huygens Essential software (version 19.04, Scientific Volume Imaging, The Netherlands; http://svi.nl). The reagents are listed in the **Supplementary Table 6.**

### FISH analysis

Image analysis was performed using a custom-written plugin (https://github.com/orion-cirb/Astro_Dots) developed for the Fiji software [42], incorporating functionalities from the Bio-Formats [43], CLIJ [44], 3D ImageJ Suite [45], and Local Thickness (https://imagej.net/plugins/local-thickness) libraries. This tool is an updated version of the previously described AstroDot plugin [22]. ROIs enclosing individual astrocytes were manually drawn using the polygon selection tool on the maximum intensity projection of each image stack in Fiji. Astrocytes in direct contact or located at a maximum distance of 165 µm from an Aβ plaque were considered as ‘close’ to plaques. Each ROI was labeled in the ROI Manager using the format (roi_number–z_start–z_stop) and saved as .roi or .zip files for further processing. Nuclei were detected using the 2D-stitched mode of Cellpose 2 [8], with the following parameters: model = ‘cyto2’, diameter = 80 pixels, flow threshold = 0.4, cell probability threshold = 0.0, and stitching threshold = 0.75. The resulting 3D nuclei were filtered by volume, retaining only those within 50–1200 µm³ to exclude false positives. The astrocyte channel was processed using a 3D median filter (σ_xy_ = 2, σ_z_= 1) and segmented using Huang automated thresholding method. A second 3D median filter (σ_xy_= 1, σ_z_ = 1) was then applied to smooth the binary mask. RNA dots were segmented by sequentially applying *a 2D Difference* of Gaussians filter (σ_1_ = 1, σ_2_ = *2*) slice-by-slice, *followed* by the Triangle thresholding method, and a 3D median filter (σ_xy_ = 1, σ_z_ = 1). Background noise in the RNA channel was estimated as the mean intensity of voxels that did not belong to any detected dot.

Subsequent analysis was conducted individually within each ROI, over the z-range defined in its name (z_start to z_stop), corresponding to a single astrocyte. The astrocyte nucleus was selected as the largest segmented nucleus within the ROI, with a dialog box allowing the user to confirm or correct this automatic selection. To approximate the astrocyte soma, the selected nucleus was isotopically dilated by 2 µm. This dilated region was then subtracted from the binary astrocyte mask, isolating the astrocytic processes. Local thickness analysis was performed on this mask to estimate the diameter distribution of the processes. RNA dots were classified into three spatial categories: within the astrocyte soma, within astrocytic processes, and outside the astrocyte. For each compartment, the total volume of RNA dots, the estimated number of dots (assuming a mean volume of 0.003 µm³ per dot), and the background-corrected integrated intensity were calculated and saved.

### RT-qPCR

RNA was extracted from whole hippocampus or synaptogliosome (one animal per sample and 2 hippocampus per sample. N = 4 for each genotype) or purified microvessels (3 animals per sample and 6 hippocampus per sample. N = 3 for each genotype) using the Rneasy Lipid tissue mini kit (Qiagen, Hilden, Germany). cDNA was then generated using the Superscript™ III Reverse Transcriptase kit (Thermofisher). Differential cDNA levels were measured using the droplet digital PCR (ddPCR) system (Biorad) and TaqMan® copy number assay probes. Briefly, cDNA and 6-carboxyfluorescein (FAM) probes and primers (**Supplementary Table 6**) were distributed into approximately 10,000 to 20,000 droplets. Nucleic acids were then PCR-amplified in a thermal cycler (following the manufacturer conditions) and read as the number of positive and negative droplets with a QX200 Droplet Digital PCR System. The ratio for each tested gene was normalized by the total number of positive droplets against *Gapdh* RNA.

### Biochemistry and *Simple Western*

Protein extracts from hippocampal synaptogliosomes (2 hippocampi of one mouse per sample) were extracted in RIPA buffer (Sigma Aldrich) with complete protease inhibitor cocktail (Roche). Supernatants were collected by centrifugation at 10000 g during 10 min at 4°C. Protein concentration was titrated with BCA kit from Sigma (BCA-1KT). Western blot was performed using the *Simple Western Jess* technique (Bio-Techne, San Jose USA), a highly reproducible, automated, sensitive and quantitative system of capillary electrophoresis in sodium dodecyl sulfate, which combines separation of proteins by molecular weight and immunodetection. Briefly, diluted protein lysate was mixed with fluorescent master mix. 3 µL of protein mix, total protein normalization reagent, blocking reagent, wash buffer, Serpin A3N primary antibody at 2 µg/mL, secondary-HRP (ready to use “detection module”, DM-001) and chemiluminescent substrate were dispensed into designated wells in a manufacturer provided microplate. The plate was loaded into the instrument and protein was drawn into individual capillaries on a 25-capillary cassette (12-230 kDa) provided by the manufacturer (SM-SW001, Bio-Techne, San Jose USA). Raw data were generated by Compass software (Bio-Techne, San Jose USA).

### Statistics

All statistical analyses performed are listed in **Supplementary Table1-5** and can also be found in the figure legends. For analyses on mouse tissues and astrocyte cultures, we used linear mixed-effects [46]. Treatment and/or genotype were included as fixed effects, while culture or mouse were included as random effects, depending on the experimental setup. Normality of model residuals was assessed using the Shapiro–Wilk test following data transformation (Tukey transformation or log transformation, as appropriate). Homogeneity of variances between groups was evaluated using the Levene’s test. Post hoc multiple comparisons were performed using Tukey’s test when applicable. All statistical analyses were performed in *R Studio* (R software). Group differences were considered statistically significant at *p* ≤ 0.05. For qPCR analyses, differences between groups were assessed using a one-tailed Mann–Whitney test.

## Acknowledgements

We are grateful to the donors who support the charities and charitable foundations cited below and more specifically to Rachel Ajzen and Léon Iagolnitzer for their generous help. We thank the administrative platforms of the CIRB and the Collège de France for their continued support.

## References

[1] Zhang YW, Thompson R, Zhang H, Xu H. APP processing in Alzheimer’s disease. Mol Brain. 2011;4:3.

[2] Iliyasu MO, Musa SA, Oladele SB, Iliya AI. Amyloid-beta aggregation implicates multiple pathways in Alzheimer’s disease: Understanding the mechanisms. Front Neurosci. 2023;17:1081938.

[3] Thal DR, Rub U, Schultz C, Sassin I, Ghebremedhin E, Del Tredici K, et al. Sequence of Abeta-protein deposition in the human medial temporal lobe. J Neuropathol Exp Neurol. 2000;59:733–48.

[4] Porsteinsson AP, Isaacson RS, Knox S, Sabbagh MN, Rubino I. Diagnosis of Early Alzheimer’s Disease: Clinical Practice in 2021. J Prev Alzheimers Dis. 2021;8:371–86.

[5] DeTure MA, Dickson DW. The neuropathological diagnosis of Alzheimer’s disease. Mol Neurodegener. 2019;14:32.

[6] Targa Dias Anastacio H, Matosin N, Ooi L. Neuronal hyperexcitability in Alzheimer’s disease: what are the drivers behind this aberrant phenotype? Transl Psychiatry. 2022;12:257.

[7] Shankar GM, Walsh DM. Alzheimer’s disease: synaptic dysfunction and Abeta. Mol Neurodegener. 2009;4:48.

[8] De Strooper B, Karran E. The Cellular Phase of Alzheimer’s Disease. Cell. 2016;164:603–15.

[9] Escartin C, Galea E, Lakatos A, O’Callaghan JP, Petzold GC, Serrano-Pozo A, et al. Reactive astrocyte nomenclature, definitions, and future directions. Nat Neurosci. 2021;24:312–25.

[10] Abbrescia P, Signorile G, Valente O, Palazzo C, Cibelli A, Nicchia GP, et al. Crucial role of Aquaporin-4 extended isoform in brain water Homeostasis and Amyloid-beta clearance: implications for Edema and neurodegenerative diseases. Acta Neuropathol Commun. 2024;12:159.

[11] Iliff JJ, Wang M, Liao Y, Plogg BA, Peng W, Gundersen GA, et al. A paravascular pathway facilitates CSF flow through the brain parenchyma and the clearance of interstitial solutes, including amyloid beta. Sci Transl Med. 2012;4:147ra11.

[12] Rasmussen MK, Mestre H, Nedergaard M. The glymphatic pathway in neurological disorders. Lancet Neurol. 2018;17:1016–24.

[13] Bosson A, Paumier A, Boisseau S, Jacquier-Sarlin M, Buisson A, Albrieux M. TRPA1 channels promote astrocytic Ca(2+) hyperactivity and synaptic dysfunction mediated by oligomeric forms of amyloid-beta peptide. Mol Neurodegener. 2017;12:53.

[14] Paumier A, Boisseau S, Jacquier-Sarlin M, Pernet-Gallay K, Buisson A, Albrieux M. Astrocyte-neuron interplay is critical for Alzheimer’s disease pathogenesis and is rescued by TRPA1 channel blockade. Brain. 2022;145:388–405.

[15] Shah D, Gsell W, Wahis J, Luckett ES, Jamoulle T, Vermaercke B, et al. Astrocyte calcium dysfunction causes early network hyperactivity in Alzheimer’s disease. Cell Rep. 2022;40:111280.

[16] Boulay AC, Saubaméa B, Adam N, Chasseigneaux S, Mazaré N, Gilbert A, et al. Translation in astrocyte distal processes sets molecular heterogeneity at the gliovascular interface. Cell Discovery. 2017;3:17005.

[17] Sakers K, Lake AM, Khazanchi R, Ouwenga R, Vasek MJ, Dani A, et al. Astrocytes locally translate transcripts in their peripheral processes. Proc Natl Acad Sci U S A. 2017;114:E3830–E8.

[18] Mazare N, Oudart M, Cohen-Salmon M. Local translation in perisynaptic and perivascular astrocytic processes - a means to ensure astrocyte molecular and functional polarity? J Cell Sci. 2021;134.

[19] Mazare N, Oudart M, Moulard J, Cheung G, Tortuyaux R, Mailly P, et al. Local Translation in Perisynaptic Astrocytic Processes Is Specific and Changes after Fear Conditioning. Cell Rep. 2020;32:108076.

[20] Sapkota D, Kater MSJ, Sakers K, Nygaard KR, Liu Y, Koester SK, et al. Activity-dependent translation dynamically alters the proteome of the perisynaptic astrocyte process. Cell Rep. 2022;41:111474.

[21] Shim B, Ciryam P, Tosun C, Serra R, Tsymbalyuk N, Keledjian K, et al. RiboTag RNA Sequencing Identifies Local Translation of HSP70 in Astrocyte Endfeet After Cerebral Ischemia. Int J Mol Sci. 2025;26.

[22] Oudart M, Tortuyaux R, Mailly P, Mazare N, Boulay AC, Cohen-Salmon M. AstroDot - a new method for studying the spatial distribution of mRNA in astrocytes. J Cell Sci. 2020;133.

[23] Tortuyaux R, Avila-Gutierrez K, Oudart M, Mazare N, Mailly P, Deschemin JC, et al. Physiopathological changes of ferritin mRNA density and distribution in hippocampal astrocytes in the mouse brain. J Neurochem. 2023;164:847–57.

[24] Jankowsky JL, Fadale DJ, Anderson J, Xu GM, Gonzales V, Jenkins NA, et al. Mutant presenilins specifically elevate the levels of the 42 residue beta-amyloid peptide in vivo: evidence for augmentation of a 42-specific gamma secretase. Hum Mol Genet. 2004;13:159–70.

[25] Luchena C, Zuazo-Ibarra J, Valero J, Matute C, Alberdi E, Capetillo-Zarate E. A Neuron, Microglia, and Astrocyte Triple Co-culture Model to Study Alzheimer’s Disease. Front Aging Neurosci. 2022;14:844534.

[26] Behrendt G, Baer K, Buffo A, Curtis MA, Faull RL, Rees MI, et al. Dynamic changes in myelin aberrations and oligodendrocyte generation in chronic amyloidosis in mice and men. Glia. 2013;61:273–86.

[27] Quintela-Lopez T, Ortiz-Sanz C, Serrano-Regal MP, Gaminde-Blasco A, Valero J, Baleriola J, et al. Abeta oligomers promote oligodendrocyte differentiation and maturation via integrin beta1 and Fyn kinase signaling. Cell Death Dis. 2019;10:445.

[28] Lee Y, Messing A, Su M, Brenner M. GFAP promoter elements required for region-specific and astrocyte-specific expression. Glia. 2008;56:481–93.

[29] Ceyzeriat K, Ben Haim L, Denizot A, Pommier D, Matos M, Guillemaud O, et al. Modulation of astrocyte reactivity improves functional deficits in mouse models of Alzheimer’s disease. Acta Neuropathol Commun. 2018;6:104.

[30] Fol R, Braudeau J, Ludewig S, Abel T, Weyer SW, Roederer JP, et al. Viral gene transfer of APPsalpha rescues synaptic failure in an Alzheimer’s disease mouse model. Acta Neuropathol. 2016;131:247–66.

[31] Boulay AC, Saubamea B, Decleves X, Cohen-Salmon M. Purification of Mouse Brain Vessels. J Vis Exp. 2015;105.

[32] Heiman M, Kulicke R, Fenster RJ, Greengard P, Heintz N. Cell type-specific mRNA purification by translating ribosome affinity purification (TRAP). Nat Protoc. 2014;9:1282–91.

[33] Mazare N, Oudart M, Cheung G, Boulay AC, Cohen-Salmon M. Immunoprecipitation of Ribosome-Bound mRNAs from Astrocytic Perisynaptic Processes of the Mouse Hippocampus. STAR Protoc. 2020;1:100198.

[34] Dobin A, Davis CA, Schlesinger F, Drenkow J, Zaleski C, Jha S, et al. STAR: ultrafast universal RNA-seq aligner. Bioinformatics. 2013;29:15–21.

[35] Paillet J, Plantureux C, Levesque S, Le Naour J, Stoll G, Sauvat A, et al. Autoimmunity affecting the biliary tract fuels the immunosurveillance of cholangiocarcinoma. J Exp Med. 2021;218.

[36] Love MI, Huber W, Anders S. Moderated estimation of fold change and dispersion for RNA-seq data with DESeq2. Genome Biol. 2014;15:550.

[37] Subramanian A, Tamayo P, Mootha VK, Mukherjee S, Ebert BL, Gillette MA, et al. Gene set enrichment analysis: a knowledge-based approach for interpreting genome-wide expression profiles. Proc Natl Acad Sci U S A. 2005;102:15545–50.

[38] Wang J, Vasaikar S, Shi Z, Greer M, Zhang B. WebGestalt 2017: a more comprehensive, powerful, flexible and interactive gene set enrichment analysis toolkit. Nucleic Acids Res. 2017;45:W130–W7.

[39] Hanzelmann S, Castelo R, Guinney J. GSVA: gene set variation analysis for microarray and RNA-seq data. BMC Bioinformatics. 2013;14:7.

[40] Avila-Gutierrez K, Monnet H, Mailly P, Boulay AC, Cohen Salmon M. Postnatal development of perivascular astrocytic processes and detection of local mRNA and translation Methods in Molecular Biology. 2025.

[41] Avila-Gutierrez K, Slaoui L, Alvear-Perez R, Kozlowski E, Oudart M, Augustin E, et al. Dynamic local mRNA localization and translation occurs during the postnatal molecular maturation of perivascular astrocytic processes. Glia. 2024;72:777–93.

[42] Schindelin J, Arganda-Carreras I, Frise E, Kaynig V, Longair M, Pietzsch T, et al. Fiji: an open-source platform for biological-image analysis. Nat Methods. 2012;9:676–82.

[43] Linkert M, Rueden CT, Allan C, Burel JM, Moore W, Patterson A, et al. Metadata matters: access to image data in the real world. J Cell Biol. 2010;189:777–82.

[44] Haase R, Royer LA, Steinbach P, Schmidt D, Dibrov A, Schmidt U, et al. CLIJ: GPU-accelerated image processing for everyone. Nat Methods. 2020;17:5–6.

[45] Ollion J, Cochennec J, Loll F, Escude C, Boudier T. TANGO: a generic tool for high-throughput 3D image analysis for studying nuclear organization. Bioinformatics. 2013;29:1840–1.

[46] Yu Z, Guindani M, Grieco SF, Chen L, Holmes TC, Xu X. Beyond t test and ANOVA: applications of mixed-effects models for more rigorous statistical analysis in neuroscience research. Neuron. 2022;110:21–35.

[47] Baleriola J, Walker CA, Jean YY, Crary JF, Troy CM, Nagy PL, et al. Axonally synthesized ATF4 transmits a neurodegenerative signal across brain regions. Cell. 2014;158:1159–72.

[48] Walker CA, Randolph LK, Matute C, Alberdi E, Baleriola J, Hengst U. Abeta1-42 triggers the generation of a retrograde signaling complex from sentinel mRNAs in axons. EMBO Rep. 2018;19.

[49] Gamarra M, de la Cruz A, Blanco-Urrejola M, Baleriola J. Local Translation in Nervous System Pathologies. Front Integr Neurosci. 2021;15:689208.

[50] Pestka S. Inhibitors of ribosome functions. Annu Rev Microbiol. 1971;25:487–562.

[51] Pestka S, Brot N. Studies on the formation of transfer ribonucleic acid-ribosome complexes. IV. Effect of antibiotics on steps of bacterial protein synthesis: some new ribosomal inhibitors of translocation. J Biol Chem. 1971;246:7715–22.

[52] Ruan L, Kang Z, Pei G, Le Y. Amyloid deposition and inflammation in APPswe/PS1dE9 mouse model of Alzheimer’s disease. Curr Alzheimer Res. 2009;6:531–40.

[53] Carney KE, Milanese M, van Nierop P, Li KW, Oliet SH, Smit AB, et al. Proteomic analysis of gliosomes from mouse brain: identification and investigation of glial membrane proteins. J Proteome Res. 2014;13:5918–27.

[54] Vanlandewijck M, He L, Mae MA, Andrae J, Ando K, Del Gaudio F, et al. A molecular atlas of cell types and zonation in the brain vasculature. Nature. 2018;554:475–80.

[55] Zhang Y, Chen K, Sloan SA, Bennett ML, Scholze AR, O’Keeffe S, et al. An RNA-sequencing transcriptome and splicing database of glia, neurons, and vascular cells of the cerebral cortex. J Neurosci. 2014;34:11929–47.

[56] Hur JY, Frost GR, Wu X, Crump C, Pan SJ, Wong E, et al. The innate immunity protein IFITM3 modulates gamma-secretase in Alzheimer’s disease. Nature. 2020;586:735–40.

[57] Chung W-S, Clarke LE, Wang GX, Stafford BK, Sher A, Chakraborty C, et al. Astrocytes mediate synapse elimination through MEGF10 and MERTK pathways. Nature. 2013;504:394–400.

[58] de Vet HC, Kessels AG, Leffers P, Knipschild PG. A randomized trial about the perceived informativeness of new empirical evidence. Does beta-carotene prevent (cervical) cancer? J Clin Epidemiol. 1993;46:509–17.

[59] Hashimoto T, Fujii D, Naka Y, Kashiwagi-Hakozaki M, Matsuo Y, Matsuura Y, et al. Collagenous Alzheimer amyloid plaque component impacts on the compaction of amyloid-beta plaques. Acta Neuropathol Commun. 2020;8:212.

[60] Soderberg L, Kakuyama H, Moller A, Ito A, Winblad B, Tjernberg LO, et al. Characterization of the Alzheimer’s disease-associated CLAC protein and identification of an amyloid beta-peptide-binding site. J Biol Chem. 2005;280:1007–15.

[61] Thuerauf DJ, Marcinko M, Belmont PJ, Glembotski CC. Effects of the isoform-specific characteristics of ATF6 alpha and ATF6 beta on endoplasmic reticulum stress response gene expression and cell viability. J Biol Chem. 2007;282:22865–78.

[62] Thuerauf DJ, Morrison L, Glembotski CC. Opposing roles for ATF6alpha and ATF6beta in endoplasmic reticulum stress response gene induction. J Biol Chem. 2004;279:21078–84.

[63] Liu C, Zhao XM, Wang Q, Du TT, Zhang MX, Wang HZ, et al. Astrocyte-derived SerpinA3N promotes neuroinflammation and epileptic seizures by activating the NF-kappaB signaling pathway in mice with temporal lobe epilepsy. J Neuroinflammation. 2023;20:161.

[64] Han X, Lei Q, Liu H, Zhang T, Gou X. SerpinA3N Regulates the Secretory Phenotype of Mouse Senescent Astrocytes Contributing to Neurodegeneration. J Gerontol A Biol Sci Med Sci. 2024;79.

[65] Zhao N, Ren Y, Yamazaki Y, Qiao W, Li F, Felton LM, et al. Alzheimer’s Risk Factors Age, APOE Genotype, and Sex Drive Distinct Molecular Pathways. Neuron. 2020;106:727–42 e6.

[66] Zattoni M, Mearelli M, Vanni S, Colini Baldeschi A, Tran TH, Ferracin C, et al. Serpin Signatures in Prion and Alzheimer’s Diseases. Mol Neurobiol. 2022;59:3778–99.

[67] Ceyzeriat K, Abjean L, Carrillo-de Sauvage MA, Ben Haim L, Escartin C. The complex STATes of astrocyte reactivity: How are they controlled by the JAK-STAT3 pathway? Neuroscience. 2016;330:205–18.

[68] de Mezer M, Rogalinski J, Przewozny S, Chojnicki M, Niepolski L, Sobieska M, et al. SERPINA3: Stimulator or Inhibitor of Pathological Changes. Biomedicines. 2023;11.

[69] Sanfilippo C, Castrogiovanni P, Imbesi R, Vecchio M, Sortino M, Musumeci G, et al. Exploring SERPINA3 as a neuroinflammatory modulator in Alzheimer’s disease with sex and regional brain variations. Metab Brain Dis. 2025;40:83.

[70] Akbor MM, Kurosawa N, Nakayama H, Nakatani A, Tomobe K, Chiba Y, et al. Polymorphic SERPINA3 prolongs oligomeric state of amyloid beta. PLoS One. 2021;16:e0248027.

[71] Janciauskiene S, Eriksson S, Wright HT. A specific structural interaction of Alzheimer’s peptide A beta 1-42 with alpha 1-antichymotrypsin. Nat Struct Biol. 1996;3:668–71.

[72] Kershaw NJ, Murphy JM, Liau NP, Varghese LN, Laktyushin A, Whitlock EL, et al. SOCS3 binds specific receptor-JAK complexes to control cytokine signaling by direct kinase inhibition. Nat Struct Mol Biol. 2013;20:469–76.

[73] Starr R, Willson TA, Viney EM, Murray LJ, Rayner JR, Jenkins BJ, et al. A family of cytokine-inducible inhibitors of signalling. Nature. 1997;387:917–21.

[74] Dvir S, Argoetti A, Lesnik C, Roytblat M, Shriki K, Amit M, et al. Uncovering the RNA-binding protein landscape in the pluripotency network of human embryonic stem cells. Cell Rep. 2021;35:109198.

[75] Fernando CD, Jayasekara WSN, Inampudi C, Kohonen-Corish MRJ, Cooper WA, Beilharz TH, et al. A STAT3 protein complex required for mitochondrial mRNA stability and cancer. Cell Rep. 2023;42:113033.

[76] Al-Ghraiybah NF, Wang J, Alkhalifa AE, Roberts AB, Raj R, Yang E, et al. Glial Cell-Mediated Neuroinflammation in Alzheimer’s Disease. Int J Mol Sci. 2022;23.

[77] Linnerbauer M, Beyer T, Nirschl L, Farrenkopf D, Losslein L, Vandrey O, et al. PD-L1 positive astrocytes attenuate inflammatory functions of PD-1 positive microglia in models of autoimmune neuroinflammation. Nat Commun. 2023;14:5555.

[78] Kummer MP, Ising C, Kummer C, Sarlus H, Griep A, Vieira-Saecker A, et al. Microglial PD-1 stimulation by astrocytic PD-L1 suppresses neuroinflammation and Alzheimer’s disease pathology. EMBO J. 2021;40:e108662.

