## Supplementary figures and legends for "Local translation controls early reactive changes in perisynaptic astrocyte processes at pre-symptomatic stages of Alzheimer’s disease"

**Supplementary text and figures**

**Supplementary Table 1. RNAseq comparison of WT PAPs versus whole astrocytes**

**Supplementary Table 2. RNAseq comparison of PAPs APP versus WT**

**Supplementary Table 3. RNAseq comparison of APP PAPs versus whole astrocytes**

**Supplementary Table 4. RNAseq raw data**

**Supplementary Table 5. Raw data and statistics**

**Supplementary Table 6. Reagents**

**A.**

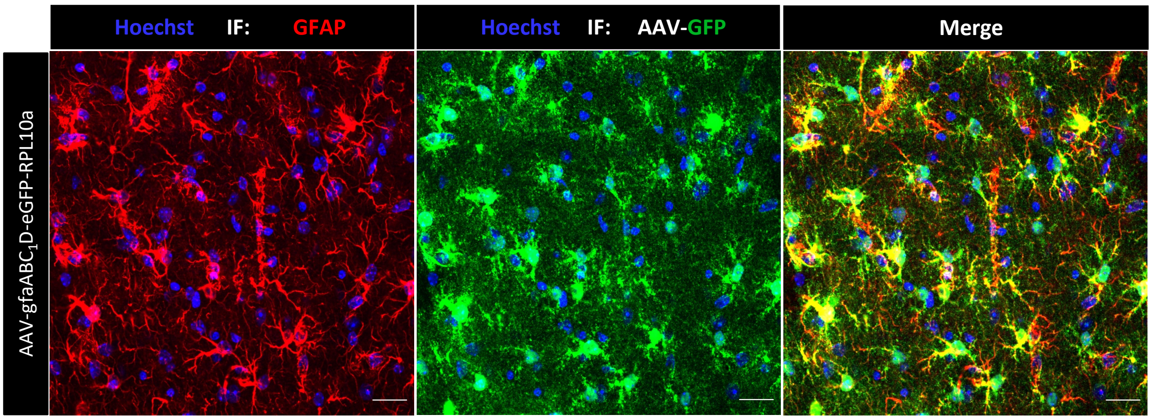


**B.**

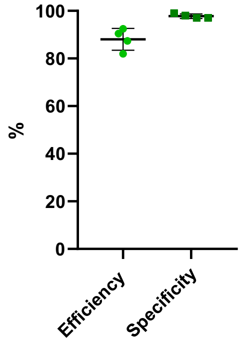


**Supplementary Fig. 1. AAV-gfaAPB1D-eGFP-Rpl10a transfection efficacy and specificity. (A)** Confocal microscopy images of hippocampal sections, stained for GFP (green) and GFAP (red) in mice injected with AAV-gfaABC$_1$D- eGFP-Rpl10a. Nuclei were counterstained with Hoechst (blue). Scale bar: 20 µm. (**B)** Quantification of AAV efficacy (% GFP$^+$GFAP$^+$ cells/number of GFAP$^+$ cells). Quantification of astrocyte specificity of AAV infection (% GFP$^+$GFAP$^+$cells/ Number of GFP$^+$ cells). N= 3-4 mice per condition. The raw data are presented in **Supplementary Table 5**.

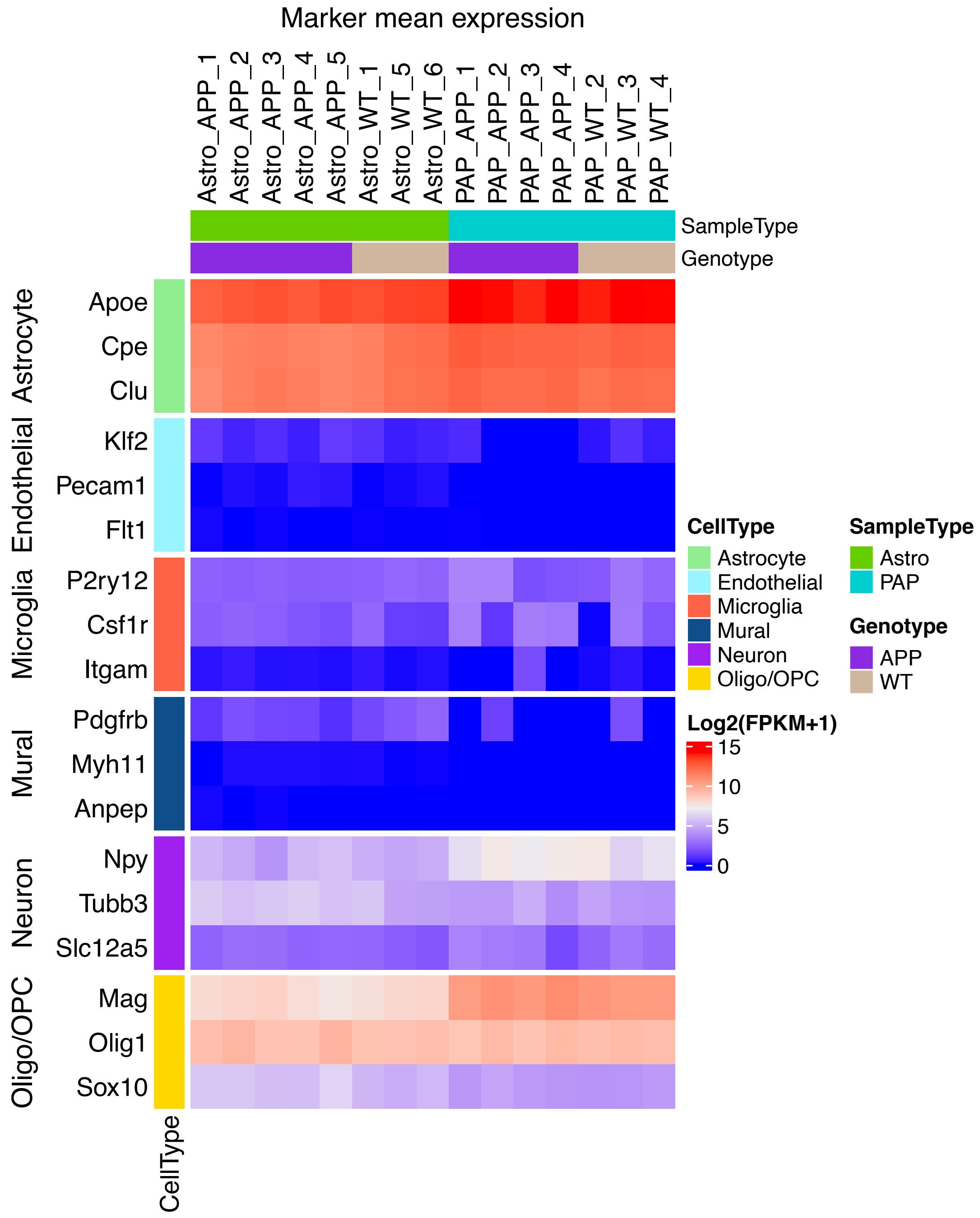


**Supplementary Fig. 2 Cell specificity of ribosome-bound mRNAs extracted by TRAP**. Enrichment genes specifically expressed in brain cells according to a previous brain single cell analysis [54, 55].

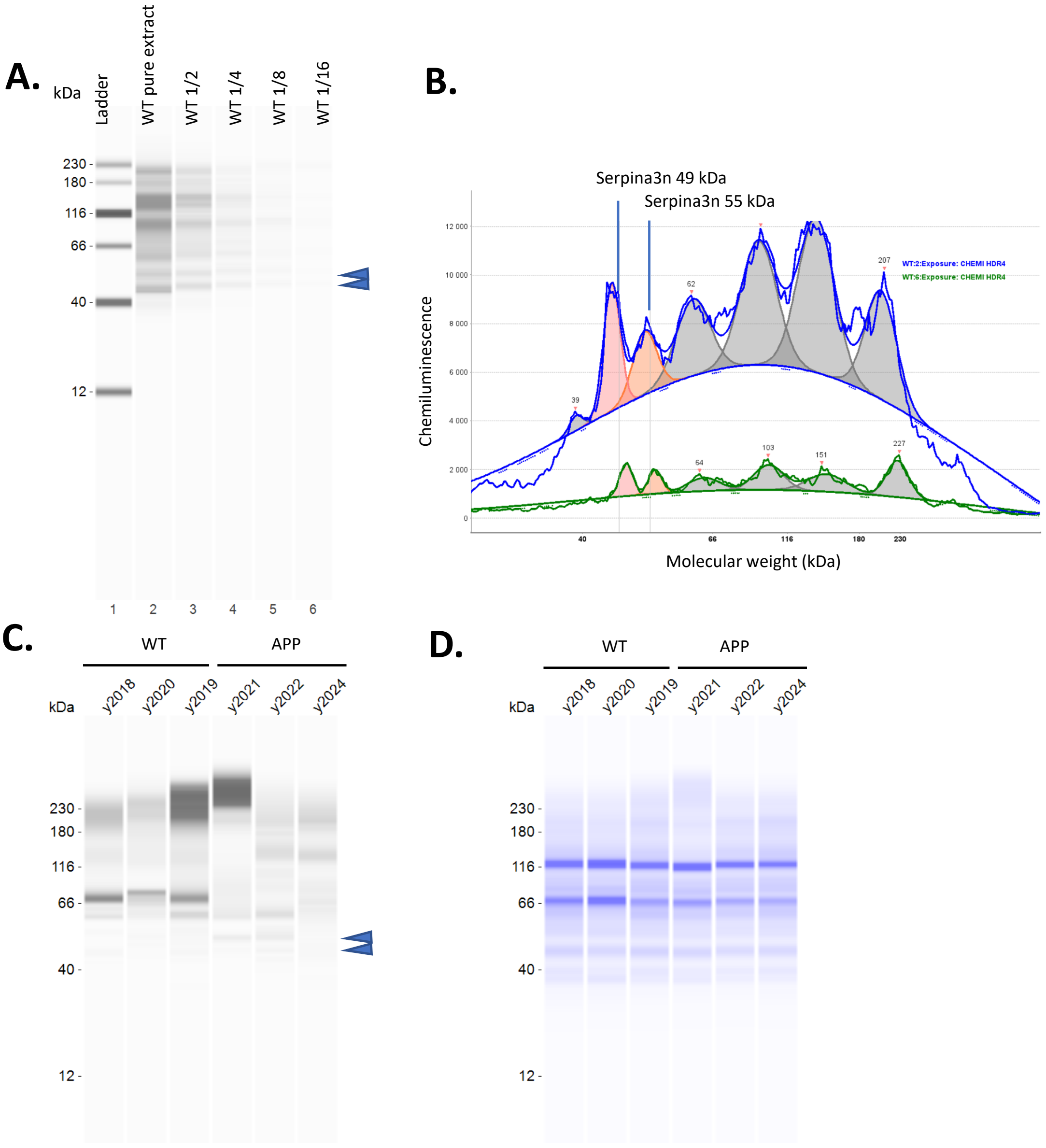


**Supplementary Fig. 3: Detection of Serpina3n in synaptogliosome protein extracts by capillary electrophoresis. A. B.** Test of antibody specificity. **A** Western blot interpretation of signals detected with increasing dilutions of the same synaptogliosome protein extract (protein prepared from the 2 hippocampi of one WT 5.5-month-old mouse; pure extract: 6 µg). Blue arrowheads show bands with the expected molecular weight for Serpina3n (50-55 kDa). **B**, Peakc analysis of the bands from lanes 2 (WT pure, in blue) and 6 (WT 1/16 dilution, in green). Peaks in pink correspond to the expected molecular weight for Serpina3n (50-55 kDa). **C.** Western blot interpretation of signals detected in 3 independent 5.5-month WT and APP synaptogliosome protein extracts (each sample corresponds to synaptogliosome extracted from the 2 hippocampi of one mouse). Blue arrowheads show bands with the expected molecular weight for Serpina3n (50-55 kDa). **D.** Coomassie blue image of the protein migration in each capillary.

**A.**

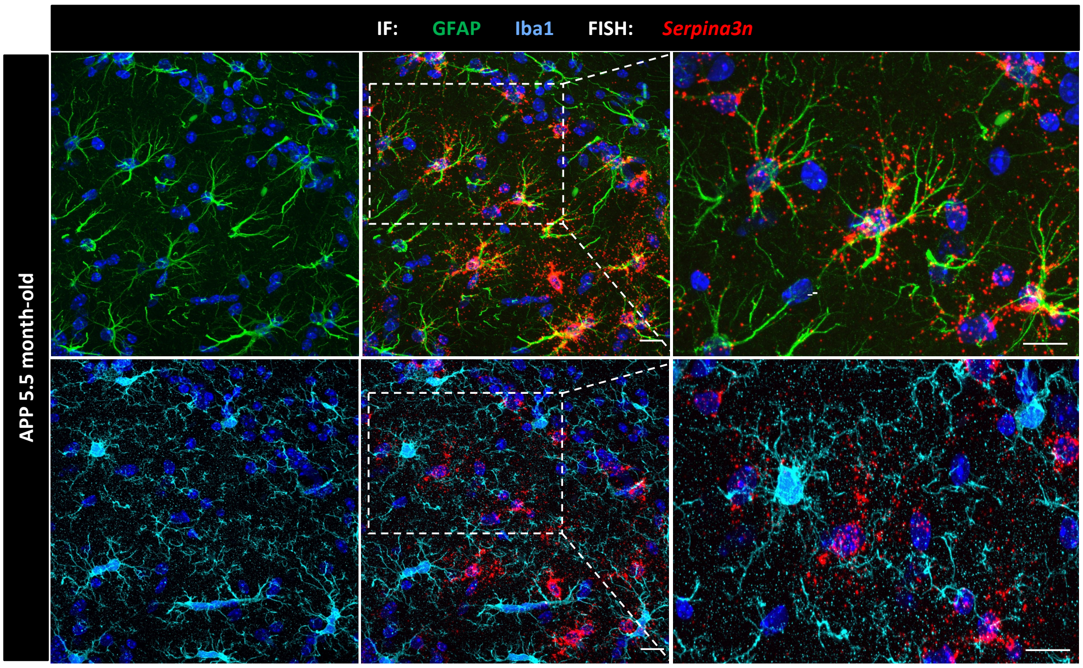


**B.**

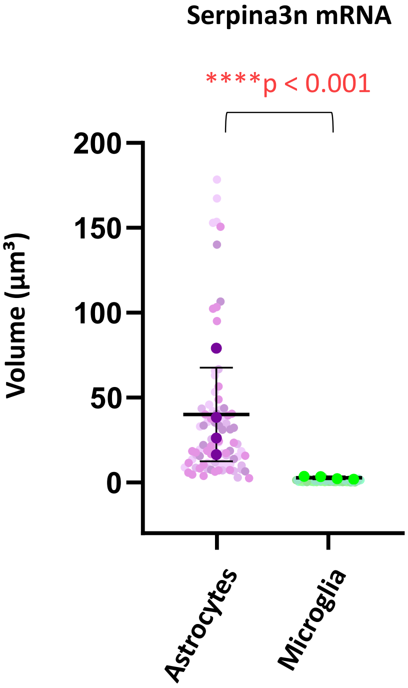

**Supplementary Fig. 4: *Serpina3n* mRNAs are not detected in microglia. (A)** Representative confocal images of a 5.5-month-old APP mouse at showing *Serpina3n* mRNAs detected by FISH (red) and Iba1 detected by immunolabeling (cyan) in CA1. Amyloid plaques are labeled by MOX4 (blue). (B) FISH quantification in CA1 hippocampal astrocytes from WT and APP/PS1 mice. Enlarged views of dotted areas in middle images are presented on the right. (**B**) Total *Serpina3n* mRNA density in astrocytes soma and microglia soma. FISH dot density was normalized to individual ROI volumes. Data are presented as mean ± SD. N = 4 mice (4 images per mouse, 70 individual astrocytes and 49 microglia). Data were analyzed using a linear mixed-effects model with **genotype as a fixed effect** and **mouse as a random effect** on Tukey transformed data. Raw data are provided in **Supplementary Table 5**

**A.**

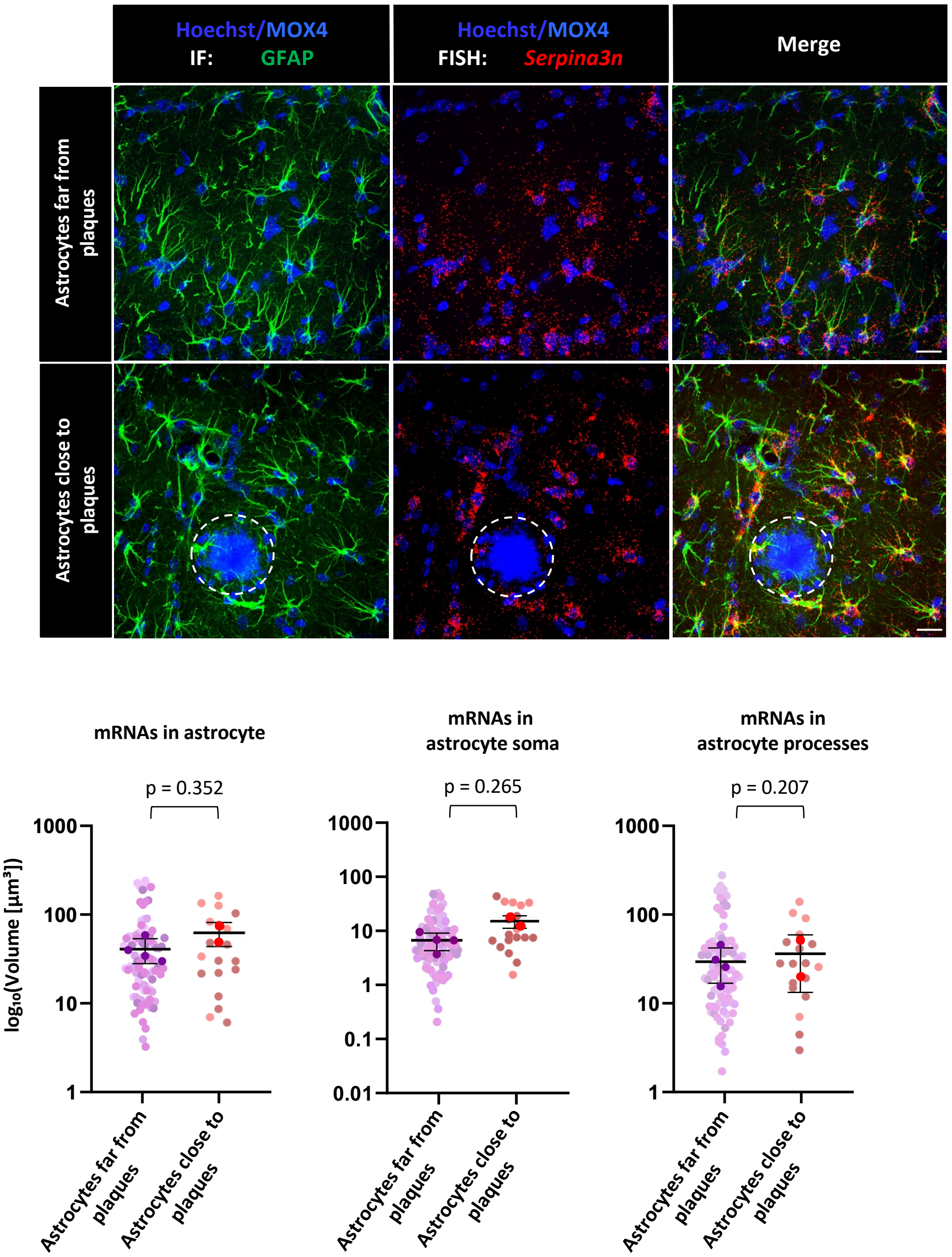


**Supplementary Fig. 5: *In situ* detection of *Serpina3n* mRNAs in 5.5 months APP in astrocytes depending on the distance with amyloid plaques. (A)** Deconvoluted confocal microscopy images showing FISH detection of *Serpina3n* mRNAs (red dots) in hippocampal astrocytes immunolabeled with GFAP (green). Amyloid-β (Aβ) plaques were stained with MOX4, and nuclei were counterstained with Hoechst (blue). Dotted circles outline Aβ plaques. Scale bar: 60 µm. (**B)** FISH quantification in CA1 hippocampal astrocytes from WT and APP/PS1 mice. Astrocytes in direct contact or located at a maximum distance of 165 µm from an Aβ plaque were considered as "close" to plaques. Total *Serpina3n* mRNA density in whole astrocytes, soma, and processes. FISH dot density was normalized to individual ROI volumes. Data are presented as mean ± SD. N = 4 WT mice and 4 APP mice (only 2 had few amyloid plaques on the 4 analyzed). Raw data are provided in **Supplementary Table 5.** Scale bar: 60 µm.

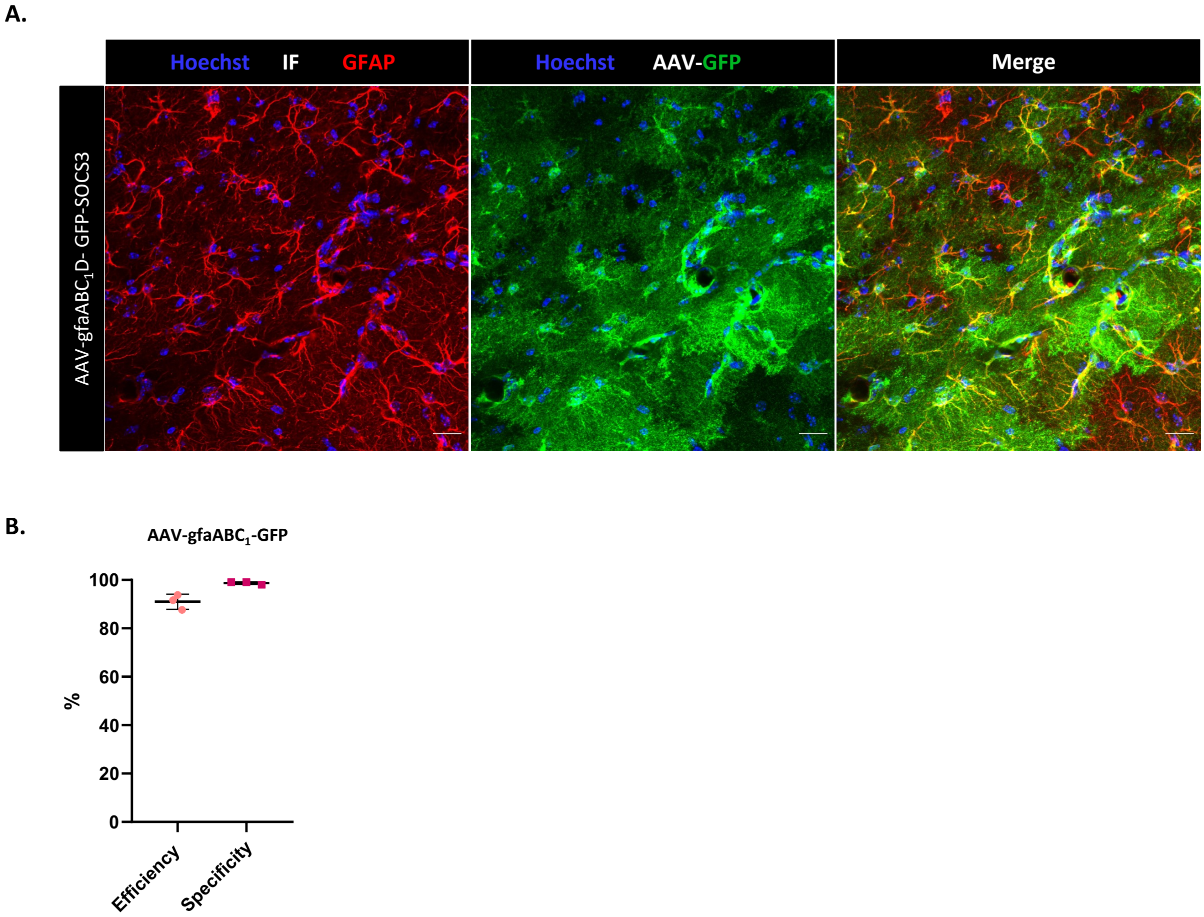


**Supplementary Fig. 6**: **AAV transfection efficacy and specificity. (A)** Confocal microscopy images of hippocampal sections, stained for GFP (green) and GFAP (red) in mice injected with AAV-gfaABC$_1$D-SOCS3 and AAV-gfaABC$_1$D-eGFP (Fig. 6). Nuclei were counterstained with Hoechst (blue). Scale bar: 20 µm. (**B)** Quantification of AAV efficacy (% GFP$^+$GFAP$^+$ / GFAP$^+$ cells). Quantification of astrocyte specificity of AAV infection (% GFP$^+$GFAP$^+$/ GFP$^+$cells). N = 3 mice per condition. Raw data are presented in **Supplementary Table 5**.
